# SPIN1 selectively silences evolutionarily young transposable elements through chromatin regulation

**DOI:** 10.64898/2026.08.20.746118

**Authors:** Hiromi Yamada, Chikara Takeuchi, Clive Barker, Nayuta Yakushiji-Kaminatsui, Aoi Shibuya, Koshi Imami, Haruhiko Koseki, Yuka W. Iwasaki

## Abstract

Mammalian genomes encode multiple transposable element (TE) silencing pathways that distinguish their targets through different molecular features, with KRAB zinc finger proteins recognizing DNA sequence, the HUSH complex sensing intronless transcripts, and the PIWI–piRNA pathway using small RNA guides. How chromatin state itself contributes to TE recognition remains less defined. Here we identify Spindlin1 (SPIN1), a three-Tudor-domain histone reader implicated in germline piRNA-directed DNA methylation, as a transcriptional repressor of evolutionarily young TEs in mouse embryonic stem cells. SPIN1 selectively binds LINE and ERV loci carrying H3K4me3 and H3K9me3, a chromatin signature enriched at young, transcription-permissive elements, and recognition of these marks by Tudor domains 1 and 2 is required for TE targeting. SPIN1 engages SPINDOC as a Tudor 1 and 3-dependent cofactor whose loss phenocopies SPIN1 depletion, and associates with the H3K9 methyltransferases SETDB1 and G9a. SPIN1 loss reduces H3K9me3 and is accompanied by increased chromatin accessibility, without altering DNA methylation. Thus, SPIN1 uses a histone-state-based mechanism to identify and repress young TEs in pluripotent cells, mechanistically distinct from its germline mode in which SPIN1 cooperates with the PIWI–piRNA pathway to promote DNA methylation, illustrating how a single histone reader engages distinct silencing machineries across cellular contexts.

## Introduction

Transposable elements (TEs) constitute a substantial fraction of mammalian genomes and pose a continuous threat to genome integrity^1,2^. Their uncontrolled mobilization can cause insertional mutagenesis, genomic rearrangements, and dysregulation of host gene expression, contributing to infertility, developmental disorders, and disease including cancer^3,4^. To counter these threats, host organisms have evolved layered epigenetic defenses that suppress TE activity at both transcriptional and post-transcriptional levels, with mechanisms that vary across developmental stages and cell types^5–7^.

A central question of TE silencing is how silencing machineries distinguish their targets from active host genes. Several recognition modalities have been characterized to date. KRAB zinc finger proteins (KRAB-ZFPs) recognize specific DNA sequences within retroviral elements and recruit TRIM28/KAP1 to deposit H3K9me3^8–11^. The Human Silencing Hub (HUSH) complex senses long, intronless nascent transcripts that characterize TE expression and engages SETDB1 to establish H3K9me3-marked heterochromatin^12–14^. In the germline, the PIWI-interacting RNA (piRNA) pathway employs sequence-specific small RNA guides to direct nuclear PIWI proteins, which recruit cofactors including SPOCD1 to drive de novo DNA methylation through DNMT3C and DNMT3L^15–19^. Each of these pathways recognizes its targets through a distinct molecular feature: DNA sequence, transcript architecture (length and lack of introns), or RNA sequence. Whether chromatin state itself can serve as a recognition signal for selective TE silencing has remained largely unexplored.

SPIN1 (Spindlin1) is a histone reader containing three Tudor-like domains. Tudor 2 recognizes the active mark H3K4me3 through its aromatic cage, while Tudor 1 binds the asymmetric dimethylarginine mark H3R8me2a in an adjacent aromatic cage; the two pockets jointly enable combinatorial readout of the H3K4me3/R8me2a methylation pattern^20,21^. Tudor 1 can alternatively engage the repressive mark H3K9me3/2 within the same binding pocket, enabling H3K4me3/K9me3 readout^22,23^. Functionally, SPIN1 has been characterized predominantly as a transcriptional activator^20,23,24^. At ribosomal DNA (rDNA) promoters, SPIN1 binding is associated with chromatin opening and stimulates RNA polymerase I-mediated rRNA synthesis^20,23^, and SPIN1 also activates Wnt/β-catenin target genes^24^. In addition, its overexpression in human cancers correlates with enhanced proliferation^25^. In the male germline, recent work identified SPIN1 as a SPOCD1-interacting cofactor that contributes to piRNA-directed DNA methylation of TEs, where its loss results in spermatogenic failure and male infertility^16,17^. In this germline context, SPIN1 acts as part of the PIWI–piRNA pathway, with TE silencing depending on small RNA-guided targeting and DNMT3C-mediated DNA methylation as the principal repressive output. However, *Spin1* is broadly expressed across embryonic and adult tissues^26^, and *Spin1*-knockout (KO) mice show postnatal lethality with defects beyond the germline^27^, suggesting additional roles in non-germline contexts where the piRNA pathway is not operational. Whether and how SPIN1 contributes to TE control outside the germline, and whether such activity employs the same or distinct molecular mechanisms compared with its germline function, has remained unclear.

Here, we identify SPIN1 as a transcriptional repressor of evolutionarily young TEs in mouse embryonic stem cells (mESCs), where the PIWI–piRNA pathway is not operational. SPIN1 selectively binds a subset of LINE and ERV loci that carry both H3K4me3 and H3K9me3, a chromatin signature enriched at evolutionarily young, transcription-permissive elements, and we show that this histone-mark recognition by Tudor domains 1 and 2 is functionally required for SPIN1 to identify its TE targets. Quantitative mass spectrometry reveals SPINDOC as a Tudor 1 and 3-dependent SPIN1 cofactor, and SPIN1 also associates with the H3K9 methyltransferases SETDB1 and G9a. Loss of SPIN1 reduces H3K9me3 and increases chromatin accessibility at target loci without altering DNA methylation, and *Spindoc*-KO cells phenocopy the TE derepression observed upon SPIN1 loss, establishing SPINDOC as a functionally required cofactor. These findings define a piRNA-independent, chromatin-based recognition and silencing mechanism for evolutionarily young TEs in pluripotent cells, in which SPIN1 engages a distinct silencing machinery from the SPOCD1– and DNA-methylation-coupled mode it employs in the germline.

## Results

### SPIN1 represses transposons in mouse embryonic stem cells

To investigate whether SPIN1 contributes to TE silencing outside the germline, we used mouse embryonic stem cells (mESCs), where the PIWI–piRNA pathway is not operational. Because TE silencing is robust in wild-type mESCs and can mask the contribution of individual factors, we used a sensitized background lacking *Pcgf6*, a Polycomb component that restricts retroelement expression^28^, together with deletion of the 2C-program transcription factor *Dux* to circumvent the proliferative arrest associated with *Pcgf6* loss^29^ (*Pcgf6/Dux* double-KO mESCs, hereafter dKO mESCs; Suppl Fig 1A,B). dKO mESCs maintain normal proliferation while exhibiting derepression of multiple TE families, particularly LINE and IAP-family ERVs (Suppl Fig 1C), and lack functional piRNA biogenesis as confirmed by the absence of nuclear piRNA pathway components (Suppl Fig 1D). This sensitized, piRNA-free system allowed us to assess SPIN1 function in TE control without confounding contributions from the germline pathway.

We generated *Spin1*-KO cell lines in the dKO background by CRISPR/Cas9-mediated targeting (Fig 1A). RNA-seq revealed widespread upregulation of TEs in *Spin1*-KO cells, including LINE and ERV elements (Fig 1B), with quantitative RT-PCR confirming derepression of multiple representative elements (IAPLTR1, IAPEz-int, IAPEy-int, L1Md_A, and L1Md_T; Fig 1C). Importantly, this TE derepression was not restricted to the dKO background: *Spin1*-KO in unmodified wild-type mESCs also elevated TE transcripts at IAPLTR1, IAPEz-int, and IAPEy-int loci (Suppl Fig 1E), establishing that SPIN1’s silencing function is not specific to the sensitized context.

**Figure 1.**
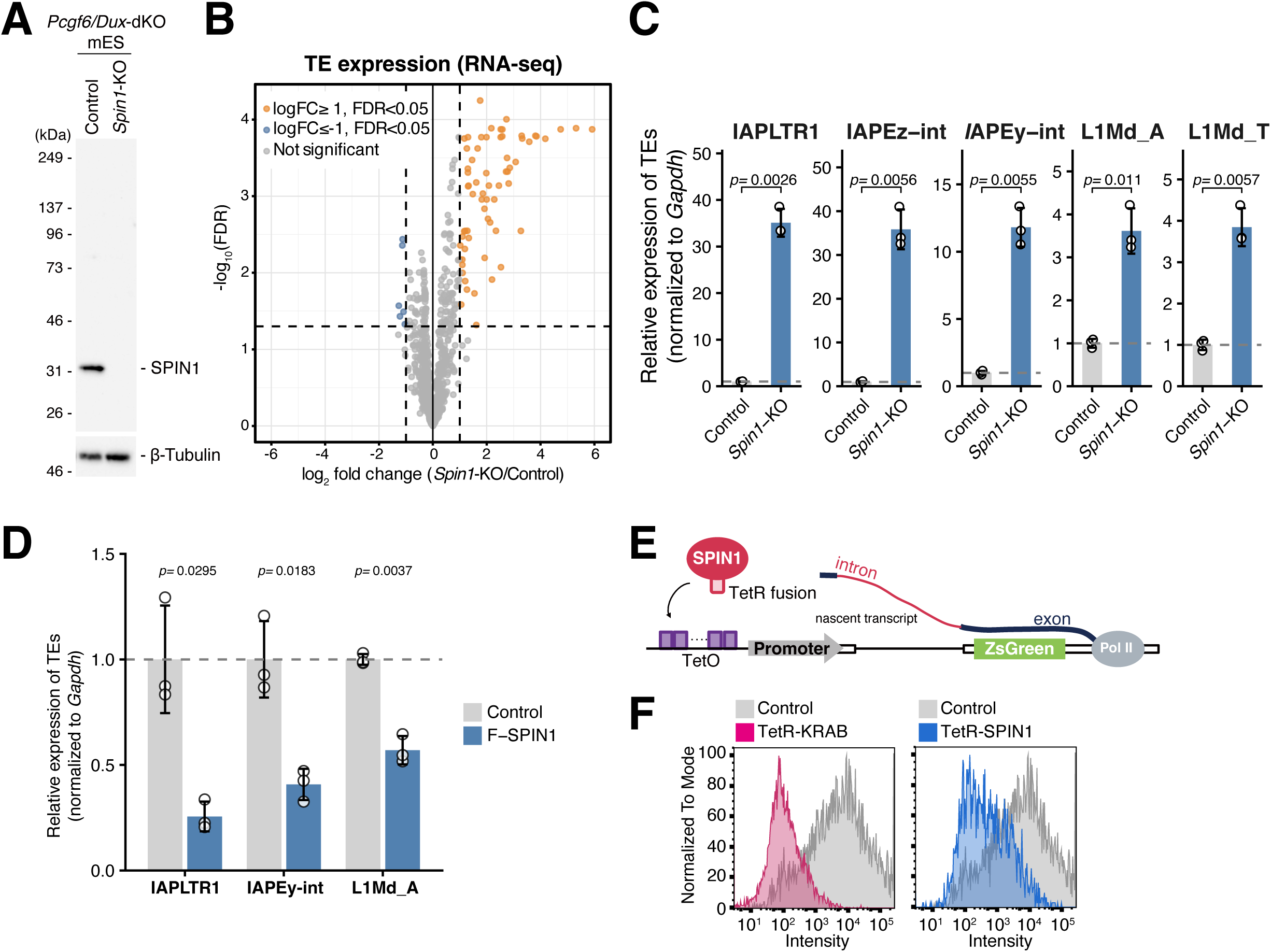
SPIN1 represses transposons in mouse embryonic stem cells. (A) Western blot analysis confirming the loss of SPIN1 protein in *Spin1*-KO cells. β-Tubulin was used as loading control. (B) Volcano plot showing differential expression of transposable elements (TEs) from RNA-seq data in *Spin1*-KO mESCs (in a *Pcgf6/Dux*-dKO background). Significantly upregulated TEs (log_2_ fold change ≥ 1, FDR < 0.05) and downregulated TEs (log_2_ fold change ≤ –1, FDR < 0.05) are highlighted in orange and blue, respectively. (C) RT-qPCR analysis of representative TEs (IAPLTR1, IAPEz-int, IAPEy-int, L1Md_A, L1Md_T) in Control and *Spin1*-KO cells. Expression levels were normalized to *Gapdh*. Data are presented as mean ± SEM from three independent experiments (*n* = 3). *P*-values were calculated using a two-sided Student’s t-test. (D) Rescue of TE silencing by re-expression of Flag-tagged wild-type SPIN1 (F-SPIN1) in *Spin1*-KO cells, as measured by RT-qPCR. Data are presented as mean ± SEM from three independent experiments (*n* = 3). *P*-values were calculated using a two-sided Student’s t-test. (E) Schematic representation of the TetR-tethering reporter system. TetR-fused candidate proteins are recruited to the *TetO* operator located upstream of an *IAP* promoter-driven *ZsGreen* reporter. (F) Flow cytometry histograms of the tethering assay in mESCs. Tethering of TetR-SPIN1 (blue) induces robust repression of the *ZsGreen* reporter compared to the TetR-only control (gray). TetR-KRAB serves as a positive control for silencing.

To test whether TE derepression reflects a SPIN1-dependent effect rather than clonal or off-target effects, we performed rescue experiments. Reintroduction of FLAG-tagged SPIN1 (Flag-SPIN1) into *Spin1*-KO cells restored silencing of derepressed TEs to levels comparable to those of control cells (Fig 1D), indicating that SPIN1 acts as a direct repressor of these elements. To further test whether direct recruitment of SPIN1 to a target locus is sufficient to drive transcriptional repression, we employed a tethering reporter assay. We constructed a reporter in which a TE-derived promoter drives expression of an intron-containing ZsGreen cassette, with upstream tetO sites allowing recruitment of TetR-fused candidate proteins to the promoter region (Fig 1E). Tethering of the well-characterized KRAB protein potently silenced reporter expression, validating the assay (Fig 1F, left). Tethering of TetR-SPIN1 produced robust reporter silencing comparable to KRAB (Fig 1F, right), demonstrating that direct recruitment of SPIN1 to a target locus is sufficient to drive transcriptional repression. These results establish SPIN1 as a transcriptional repressor of TEs in mESCs and identify it as a candidate chromatin-based silencing factor that operates outside the germline piRNA pathway.

### SPIN1 selectively binds evolutionarily young transposons marked by both H3K4me3 and H3K9me3

To define how SPIN1 identifies its TE targets, we mapped its genome-wide binding by CUT&Tag, alongside the active mark H3K4me3 and the repressive mark H3K9me3. Across the genome, the majority of H3K4me3 and H3K9me3 peaks were non-overlapping, but a discrete subset of 2,502 sites carried both modifications (Fig 2A). At these co-marked sites, SPIN1 was strongly enriched, whereas SPIN1 binding was minimal at sites carrying H3K4me3 alone or H3K9me3 alone (Fig 2B). Genomic feature annotation revealed that the H3K4me3/H3K9me3 co-marked sites were predominantly located within LINE elements (2,008 of 2,502 sites; Fig 2C), with smaller contributions from other TE or repeats and a minor fraction within genes (121 sites) or intergenic regions (27 sites). Notably, SPIN1 enrichment scaled with H3K4me3 and H3K9me3 signal intensity regardless of the underlying genomic feature class (LINE, other repeats, genes, or intergenic regions; Suppl Fig 2A), indicating that SPIN1 binding is determined by the local chromatin state rather than by the genomic feature itself.

**Figure 2.**
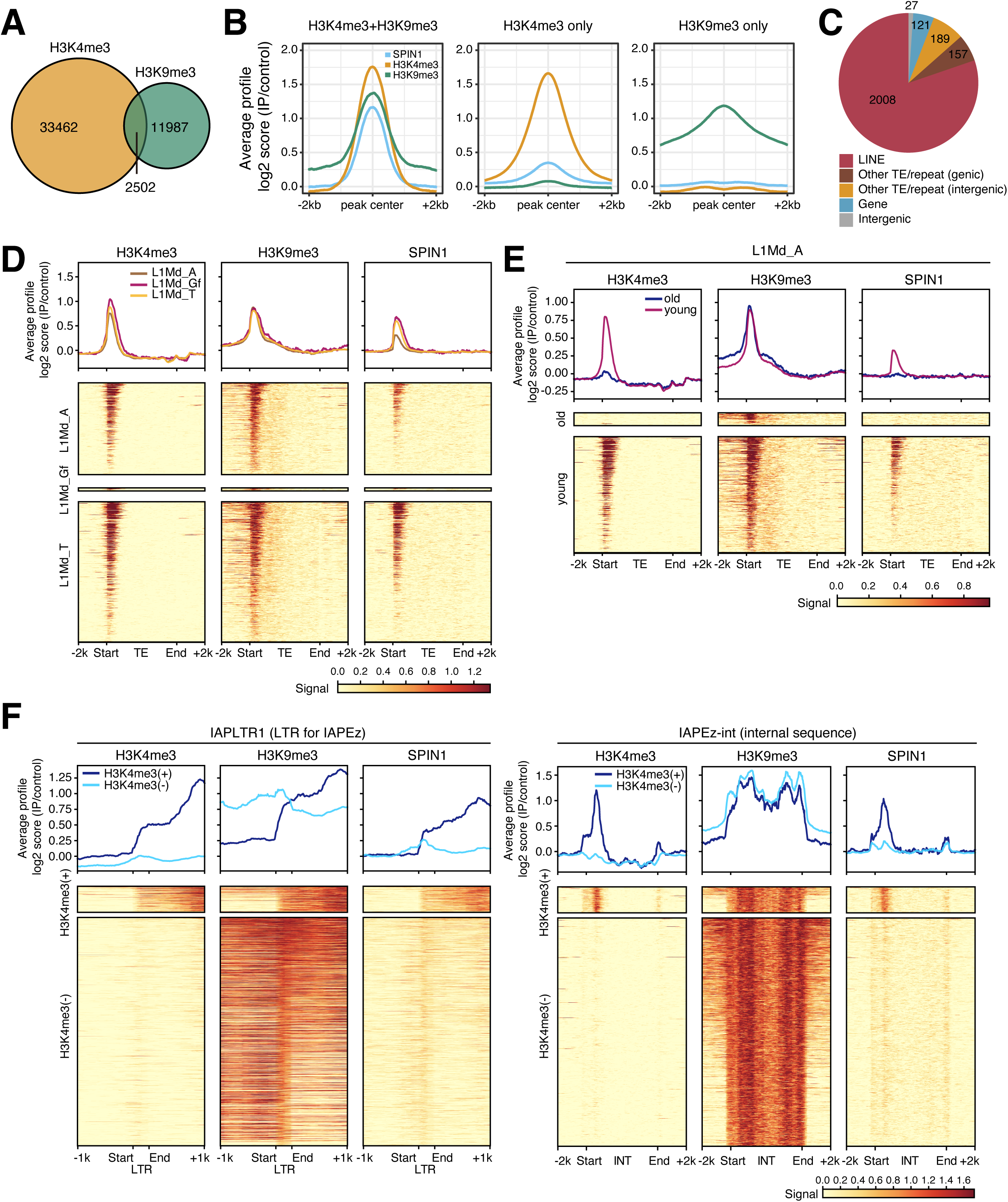
SPIN1 binds evolutionarily young transposons marked by both H3K4me3 and H3K9me3. (A) Venn diagram showing the overlap between H3K4me3 and H3K9me3 CUT&Tag peaks in mESCs. A total of 2,502 regions are marked by both modifications. (B) Average CUT&Tag profiles of SPIN1, H3K4me3, and H3K9me3 at different categories of genomic regions: regions marked by both H3K4me3 and H3K9me3 (left), H3K4me3 only (middle), and H3K9me3 only (right). SPIN1 enrichment is specifically observed at the sites with both H3K4me3 and H3K9me3 marks. (C) Pie chart showing the H3K4me3/H3K9me3 co-marked sites. The majority of SPIN1 occupancy is concentrated at LINE elements and other TE/repeat. (D) Meta-plots (top) and heatmaps (bottom) of H3K4me3, H3K9me3, and SPIN1 signals across representative LINE families (L1Md_A, L1Md_Gf, and L1Md_T). The signals are centered on the indicated TE sequences. (E) Comparison of H3K4me3, H3K9me3, and SPIN1 enrichment between evolutionarily young and old L1Md_A elements. Average profiles (top) and heatmaps (bottom) show that SPIN1 and the associated H3K4me3 and H3K9me3 marks are specifically enriched at young L1Md_A elements. (F) Meta-plots (top) and heatmaps (bottom) of H3K4me3, H3K9me3, and SPIN1 occupancy at IAP-LTR1 (LTR for IAPEz, left) and IAPEz-int (internal sequence, right). Individual TE copies were clustered based on H3K4me3 signal intensity, which separated them into H3K4me3-high (dark blue) and H3K4me3-low (light blue) clusters.

We next examined SPIN1 occupancy at evolutionarily young LINE1 families. In the mouse genome, the youngest L1 lineages comprise the L1Md (LINE-1 Mus domesticus) families, of which L1Md_A, L1Md_Gf, and L1Md_T are the most recently active and retain retrotransposition potential^30,31^. Average profiles and per-element heatmaps revealed pronounced SPIN1 binding at the 5′ end, near the transcription start site, of L1Md_A, L1Md_Gf, and L1Md_T elements, coinciding with H3K4me3 and H3K9me3 signals (Fig 2D). In contrast, SPIN1 binding was negligible at older LINE1 families such as L1Md_F or at the ERVK family MMERVK10C^30,31^. These elements carried robust H3K9me3 but lacked detectable H3K4me3 (Suppl Fig 2B), suggesting that the absence of H3K4me3, rather than H3K9me3 levels, accounts for their exclusion from SPIN1 targeting. Thus, SPIN1 does not associate uniformly with H3K9me3-marked heterochromatin but instead requires the additional presence of H3K4me3 for selective binding.

To address this directly within a single LINE1 family, we stratified L1Md_A elements into evolutionarily young and old subgroups based on sequence divergence from consensus. Young L1Md_A copies carried robust H3K4me3 together with H3K9me3 and were strongly bound by SPIN1, whereas older L1Md_A copies lacked H3K4me3, retained only weak H3K9me3, and showed no detectable SPIN1 enrichment (Fig 2E). Even within the same TE family, evolutionary age thus correlates with H3K4me3 status, which in turn predicts SPIN1 occupancy.

Whereas LINE1 families could be stratified by evolutionary age through sequence divergence, ERV elements show greater heterogeneity in copy structure and age within families, making straightforward age-based stratification less tractable. The IAP family analyzed here represents murine-specific ERVs that are among the youngest and most active mouse retrotransposons, with experimentally validated transposition activity^32,33^. We therefore stratified IAPEz elements based on H3K4me3 status itself. Within the IAPEz family, SPIN1 bound preferentially to copies carrying H3K4me3 over H3K4me3-negative copies, despite comparable H3K9me3 levels in both groups; this pattern was evident at both IAP-LTR1 sequences (the LTRs flanking IAPEz elements) and IAPEz-int internal sequences (Fig 2F). Notably, the H3K4me3-positive subset represented only a small fraction of total IAPEz elements, yet *Spin1*-KO produced substantial derepression of multiple ERV families at the transcriptome level (Suppl Fig 2C). This disproportionate impact suggests that SPIN1 acts on a small but transcriptionally active subset of ERV elements, those that carry H3K4me3 and are presumably permissive to transcription, where its silencing function is particularly critical.

These analyses identify the co-occurrence of H3K4me3 and H3K9me3 as a chromatin signature for evolutionarily young, transcriptionally permissive TEs across both LINE and ERV classes. SPIN1 occupancy correlates with this chromatin signature.

### Histone modification recognition and protein interaction by SPIN1 are required for transposon silencing

The selective binding of SPIN1 to H3K4me3/H3K9me3 co-marked young TEs raised two related questions. First, whether the recognition of these histone modifications is functionally required for SPIN1-mediated TE silencing. Second, whether the same SPIN1 molecule must recognize both modifications simultaneously, since bulk genomic profiling cannot distinguish concurrent recognition within a single molecule from independent recognition by separate SPIN1 molecules within a heterogeneous cell population. To address both questions, we dissected the contribution of individual SPIN1 domains to TE silencing.

SPIN1 contains an N-terminal intrinsically disordered region (IDR) that mediates liquid-liquid phase separation^34^, followed by three Tudor-like domains (Fig 3A). Tudor 1 recognizes the asymmetric dimethylarginine mark H3R8me2a and can also engage the repressive mark H3K9me3; Tudor 2 primarily binds the active mark H3K4me3; together, the adjacent Tudor 1 and Tudor 2 binding pockets enable combinatorial readout of these histone modifications. Tudor 3 mediates protein-protein interactions with multiple partners including SPOCD1 and SPINDOC^17,20–23,35,36^. However, whether and how these activities contribute to TE silencing remains unclear. We took advantage of this domain organization to dissect which SPIN1 activities are required for TE repression.

**Figure 3.**
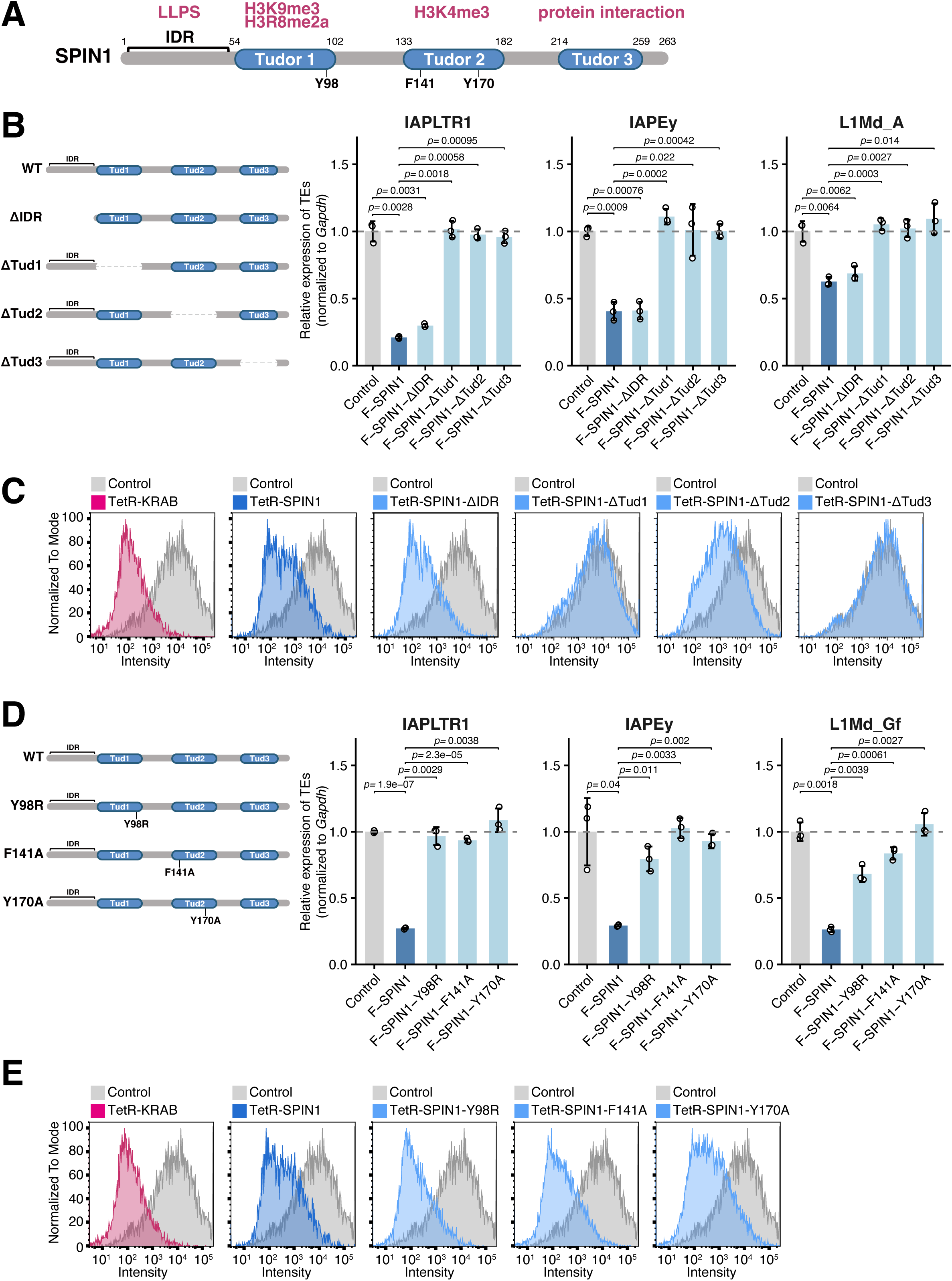
Recognition of histone modifications by SPIN1 is required for transposon silencing. (A) Schematic representation of the SPIN1 domain structure. SPIN1 consists of an N-terminal intrinsically disordered region (IDR) associated with liquid-liquid phase separation (LLPS), followed by three Tudor-like domains. The Tudor 1 domain recognizes H3K9me3 and H3R8me2a (Y98 residue), while the Tudor 2 domain recognizes H3K4me3 (F141 and Y170 residues). The Tudor 3 domain is involved in protein-protein interactions with factors such as SPOCD1 and SPINDOC. (B) Rescue of TE silencing in *Spin1*-KO mESCs by re-expression of Flag-tagged wild-type (F-SPIN1) or deletion mutants. Expression levels of the indicated TEs were measured by RT-qPCR and normalized to *Gapdh*. Full-length and IDR-deleted SPIN1 variants effectively restore silencing, whereas mutants lacking any of the Tudor domains fail to rescue TE repression. Data are presented as mean ± SEM from three independent experiments (*n* = 3). *P*-values were calculated using a two-sided Student’s t-test. (C) Flow cytometry histograms of the tethering assay in mESCs using TetR-fused SPIN1 deletion mutants. Tethering of TetR-SPIN1, SPIN1-ΔIDR, and SPIN1-ΔTud2 (light blue) induces repression of the reporter compared to the TetR-only control (gray), whereas SPIN1-ΔTud1 and SPIN1-ΔTud3 fail to repress. TetR-KRAB serves as a positive control. (D) Rescue of TE silencing by re-expression of Flag-tagged wild-type or point mutants targeting the histone-binding pockets (Y98R in Tudor 1; F141A and Y170A in Tudor 2) in *Spin1*-KO mESCs. All point mutants fail to restore TE silencing, highlighting the requirement for histone mark recognition in endogenous TE repression. Data are presented as mean ± SEM from three independent experiments (*n* = 3). *P*-values were calculated using a two-sided Student’s t-test. (E) Flow cytometry histograms of the tethering assay using TetR-fused SPIN1 point mutants. In contrast to the results in panel **D**, tethering of TetR-SPIN1-Y98R, –F141A, or –Y170A (light blue) robustly represses the reporter, suggesting that these residues are critical for recruitment but not for the intrinsic silencing capability of SPIN1. Control, TetR-KRAB, and TetR-SPIN1 (WT) data are shared with panel **C**, as the experiments were performed in parallel.

We first generated SPIN1 deletion mutants lacking individual domains (ΔIDR, ΔTud1, ΔTud2, or ΔTud3) and assessed their ability to restore TE silencing in *Spin1*-KO mESCs. Reintroduction of full-length Flag-SPIN1 (WT) restored silencing of IAPLTR1, IAPEy, and L1Md_A, and the IDR-deleted variant (ΔIDR) showed comparable rescue activity, indicating that the IDR is dispensable for TE repression in this context (Fig 3B). In contrast, deletion of any single Tudor domain (ΔTud1, ΔTud2, or ΔTud3) abolished rescue, with TE expression remaining at levels indistinguishable from the control (Fig 3B). All three Tudor domains are therefore individually required for SPIN1-mediated TE silencing in their endogenous chromatin context, whereas the IDR is dispensable.

To distinguish whether each Tudor domain contributes to target recognition or to the silencing output itself, we turned to the tethering reporter assay, in which TetR-mediated recruitment of SPIN1 to the reporter promoter bypasses the requirement for chromatin-based target identification. While tethering of TetR-SPIN1 (WT) and TetR-SPIN1-ΔIDR efficiently silenced the reporter, tethering of TetR-SPIN1-ΔTud2 also produced reporter silencing (Fig 3C). This indicates that, when SPIN1 is forcibly recruited to chromatin, the loss of Tudor 2 no longer compromises silencing. However, tethering of either TetR-SPIN1-ΔTud1 or TetR-SPIN1-ΔTud3 failed to silence the reporter (Fig 3C). Both Tudor 1 and Tudor 3 are therefore required for the silencing output beyond their role in chromatin recruitment, suggesting that these domains may mediate interactions with downstream silencing factors.

To dissect the recognition function of Tudor 1 and Tudor 2 more specifically, we generated point mutants that selectively disrupt histone-mark binding without broadly destabilizing the domains. The Y98R mutation in Tudor 1 abolishes H3K9me3 recognition while preserving overall domain integrity, and the F141A and Y170A mutations in Tudor 2 disrupt H3K4me3 binding^20–23,35^. Reintroduction of any of these point mutants (Y98R, F141A, or Y170A) into *Spin1*-KO mESCs failed to rescue silencing of IAPLTR1, IAPEy, or L1Md_Gf, with TE expression remaining elevated comparable to *Spin1*-KO control (Fig 3D). Recognition of H3K9me3 by Tudor 1 and recognition of H3K4me3 by Tudor 2 are therefore both required for SPIN1-mediated TE silencing.

Importantly, the failure of either Y98R (H3K9me3-binding deficient but H3K4me3-binding competent) or F141A and Y170A (H3K4me3-binding deficient but H3K9me3-binding competent) to rescue silencing addresses the question of whether the same SPIN1 molecule must recognize both modifications. The fact that disrupting either single recognition activity completely abolishes rescue supports a model in which both H3K4me3 and H3K9me3 recognition activities are simultaneously required for productive target selection, consistent with combinatorial readout by SPIN1’s Tudor 1 and Tudor 2 binding pockets reported in vitro^22,23^.

When tested in the tethering reporter assay, TetR-SPIN1-Y98R, TetR-SPIN1-F141A, and TetR-SPIN1-Y170A all retained silencing activity comparable to WT SPIN1 (Fig 3E). This confirms that the failure of these point mutants to rescue endogenous TE silencing reflects a specific defect in target recognition rather than a loss of intrinsic repressive activity, and it further clarifies the role of Tudor 1. Although deletion of Tudor 1 abolished tethering-mediated silencing, the point mutation that specifically disrupts H3K9me3 binding within Tudor 1 (Y98R) did not, indicating that Tudor 1 contributes to silencing through two separable activities.

These analyses establish that recognition of H3K4me3 and H3K9me3 by Tudor 1 and Tudor 2 of a single SPIN1 molecule is required for target selection, while Tudor 1 and Tudor 3 together mediate the silencing output possibly through interactions with downstream factors.

### SPINDOC is a SPIN1 cofactor required for transposon silencing

Our domain analysis revealed that Tudor 1 and Tudor 3 of SPIN1 are both required for the silencing output, even when SPIN1 is forcibly recruited to chromatin by tethering. This requirement suggested that these two domains mediate interactions with downstream silencing factors. To directly identify factors whose binding to SPIN1 depends on Tudor 1 or Tudor 3, we performed three parallel quantitative mass spectrometry (IP-MS) experiments using PA-tagged SPIN1 immunoprecipitates from mESCs. We compared full-length PA-SPIN1 immunoprecipitates against PA-mScarlet immunoprecipitates to identify general SPIN1 interactors, PA-SPIN1-ΔTud1 immunoprecipitates to identify factors requiring Tudor 1, and PA-SPIN1-ΔTud3 immunoprecipitates to identify factors requiring Tudor 3. Among the proteins significantly enriched over the mScarlet control as SPIN1-associated factors (Fig 4A), two proteins, SPINDOC and MORC4, emerged at the intersection of all three comparisons (Fig 4B; Suppl Fig 3A,B), indicating that both interactions depend on Tudor 1 and Tudor 3.

**Figure 4.**
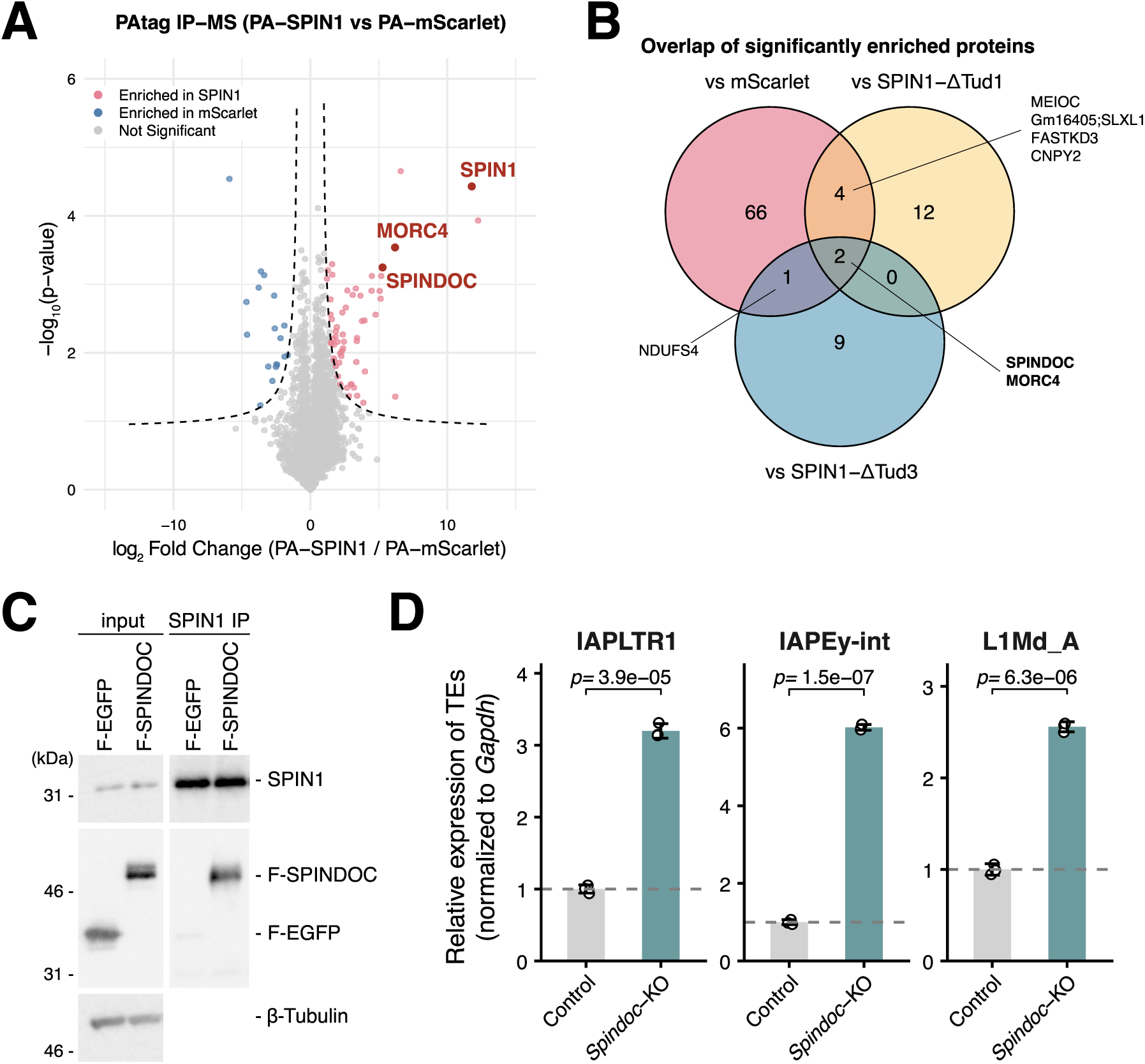
SPINDOC is a SPIN1 cofactor required for transposon silencing. (A) Volcano plot showing the enrichment of proteins co-purified with PA-tagged full-length SPIN1 (PA-SPIN1) compared to PA-mScarlet control, as determined by IP-MS (*n* = 3 biological replicates per condition). The dashed horizontal line denotes FDR = 0.05. Proteins significantly enriched in PA-SPIN1 (FDR < 0.05) are highlighted in pink. SPIN1, SPINDOC, and MORC4 are indicated (A full list of identified proteins is provided in Supplementary Table S3). (B) Venn diagram showing the overlap of proteins significantly enriched in PA-SPIN1 immunoprecipitates compared to three independent control conditions: PA-mScarlet (vs mScarlet), PA-SPIN1-ΔTud1 (vs SPIN1-ΔTud1), and PA-SPIN1-ΔTud3 (vs SPIN1-ΔTud3). The intersection of all three sets contains two proteins, SPINDOC and MORC4, indicating that their association with SPIN1 depends on both Tudor 1 and Tudor 3 domains. Selected proteins enriched in individual subsets (MEIOC, Gm16405;SLXL1, FASTKD3, CNPY2, NDUFS4) are indicated. Note that uncharacterized cluster proteins and immunoglobulin fragments were excluded from the diagram (see Methods for details). (C) Co-immunoprecipitation assay showing the interaction between endogenous SPIN1 and Flag-SPINDOC in mESCs. Flag-EGFP served as a negative control. β-Tubulin was used as a loading control for input samples. (D) RT-qPCR analysis of TE expression (IAPLTR1, IAPEy-int, L1Md_A) in Control and *Spindoc-*KO mESCs. Expression levels were normalized to *Gapdh*. Data are presented as mean ± SEM from three independent experiments (*n* = 3). P-values were calculated using a two-sided Student’s t-test.

SPINDOC is a previously characterized SPIN1-binding partner^36^. Our IP-MS analysis recovered SPINDOC as a top hit with Tudor 1 and 3 dependency, consistent with a previously characterized interaction between SPINDOC and SPIN1^36^ and revealing an additional requirement for Tudor 1 in stable cellular binding (Fig 4A and Suppl Table S3). The IP-MS analysis also identified MORC4, a member of the MORC family of chromatin-associated ATPases^37^, as an additional Tudor 1/3-dependent SPIN1 interactor that has not been previously reported as a SPIN1-binding partner. Several additional factors were identified that selectively required either Tudor 1 or Tudor 3 alone (Suppl Fig 3A,B and Suppl Table S3), but only SPINDOC and MORC4 depended on both domains (Fig 4B).

We further confirmed the SPIN1–SPINDOC interaction in mESCs by co-immunoprecipitation. F-SPINDOC, but not F-EGFP, was co-immunoprecipitated with endogenous SPIN1 (Fig 4C), confirming the physical interaction between SPIN1 and SPINDOC. To test the functional requirement of SPINDOC for SPIN1-mediated TE silencing, we generated *Spindoc*-KO mESCs and measured TE expression by RT-qPCR (Fig 4D; Suppl Fig 3C). Loss of SPINDOC resulted in robust derepression of IAPLTR1, IAPEy-int, and L1Md_A elements, recapitulating the TE derepression observed in *Spin1*-KO cells (Fig 4D). This matched phenotype, together with the identification of SPINDOC as a Tudor 1/3-dependent SPIN1-binding cofactor, indicates that SPINDOC is functionally required for SPIN1-mediated TE silencing in mESCs, consistent with our domain analysis identifying Tudor 1 and Tudor 3 as the silencing-output domains.

Together, these findings identify SPINDOC and potentially MORC4 as Tudor 1 and 3-dependent SPIN1 interactors and establish SPINDOC as a functionally required cofactor for SPIN1-mediated TE silencing in mESCs.

### SPIN1 enforces H3K9me3-based silencing without altering DNA methylation at transposons

Having identified SPIN1 and SPINDOC as a silencing axis at evolutionarily young TEs, we next asked how this complex enforces transcriptional repression at the chromatin level. SPIN1 might reinforce H3K9me3-based heterochromatin, consistent with the H3K9me3 marking observed at SPIN1 target sites. Alternatively or in addition, SPIN1 might engage the DNA methylation pathway, given its known cooperation with SPOCD1 and the DNMT3C/L machinery in the male germline^17^. To distinguish these possibilities, we profiled H3K4me3 and H3K9me3 by CUT&Tag in control and *Spin1*-KO mESCs. At SPIN1 peak centers, H3K4me3 levels were essentially unchanged upon *Spin1* loss, indicating that SPIN1 is dispensable for H3K4me3 maintenance (Fig 5A, left). In contrast, H3K9me3 was substantially reduced in *Spin1*-KO cells (Fig 5A, right), demonstrating that SPIN1 contributes to the maintenance of H3K9me3 specifically at its bound loci.

**Figure 5.**
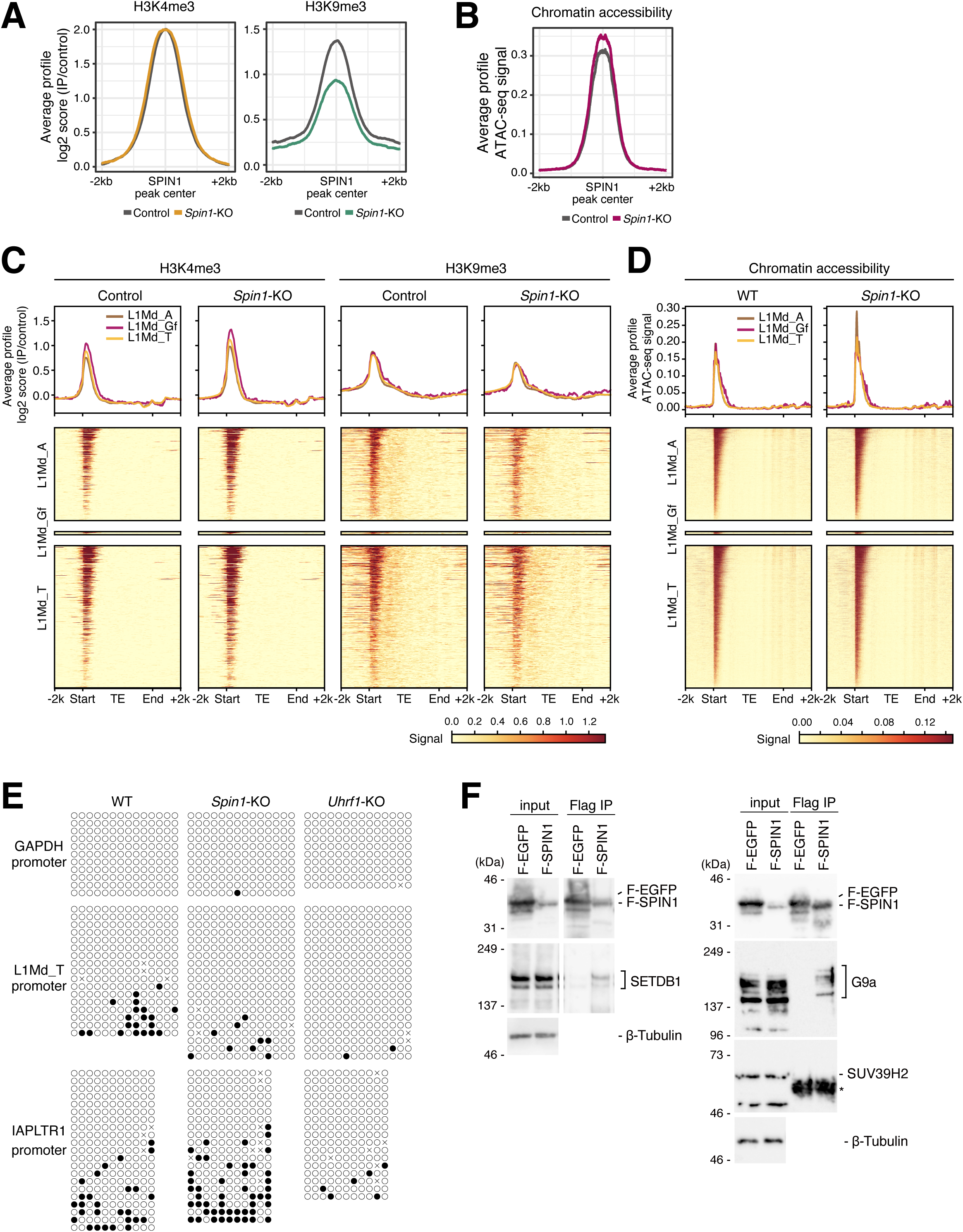
SPIN1 enforces H3K9me3-based silencing without altering DNA methylation at transposons. (A) Average CUT&Tag profiles of H3K4me3 (left) and H3K9me3 (right) at SPIN1 peak centers in Control (gray) and *Spin1-*KO cells. H3K9me3 levels are significantly reduced upon *Spin1* deletion, while H3K4me3 levels remain largely unchanged. (B) Average ATAC-seq profiles at SPIN1 peak centers, showing increased chromatin accessibility in *Spin1-*KO cells compared to control. (C) Meta-plots (top) and heatmaps (bottom) of H3K4me3 and H3K9me3 enrichment at representative L1Md families (L1Md_A, L1Md_Gf, and L1Md_T) in Control and *Spin1-* KO cells. The reduction of H3K9me3 is specifically observed at the 5′ end near the transcription start site of these elements in *Spin1-*KO cells. (D) Meta-plots (top) and heatmaps (bottom) of ATAC-seq signals across the same L1Md families (L1Md_A, L1Md_Gf, and L1Md_T), demonstrating increased accessibility at the 5′ TSS-proximal regions upon the loss of SPIN1. (E) Bisulfite sequencing analysis of DNA methylation at the *Gapdh* promoter (control), the L1Md_T promoter, and the IAPLTR1 promoter in WT, *Spin1-*KO, and *Uhrf1-*KO mESCs. Filled and open circles represent methylated and unmethylated CpG sites, respectively. *Uhrf1-*KO serves as a positive control for the loss of DNA methylation. (F) Left: Co-immunoprecipitation (Co-IP) assay in mESCs showing the interaction between Flag-SPIN1 and endogenous SETDB1. Right: Co-IP assays showing the interaction between Flag-SPIN1 and endogenous G9a, and the absence of interaction with SUV39H2. Flag-EGFP served as a negative control. The asterisk indicates signals from IgG (heavy chain). Note that the protein size of Flag-EGFP (36 kDa) and Flag-SPIN1 (35 kDa) is highly similar. β-Tubulin was used as a loading control for input samples.

To assess the functional consequence of this H3K9me3 reduction on chromatin state, we performed ATAC-seq to measure chromatin accessibility. We were particularly interested in this measurement because SPIN1 has been reported to promote chromatin opening and to facilitate transcriptional activation at ribosomal DNA promoters, where it acts as a positive regulator of RNA polymerase I-mediated rRNA synthesis^20,23^. Whether SPIN1 promotes or restricts chromatin accessibility at TE loci, where it acts as a repressor, was not predictable from these prior studies. At SPIN1 peak centers in mESCs, chromatin accessibility was increased in *Spin1*-KO cells compared with control (Fig 5B). Thus, SPIN1 acts to restrict chromatin accessibility at TE target sites, in contrast to its previously described role in promoting chromatin opening at rDNA promoters. We next examined whether these chromatin changes occurred specifically at the evolutionarily young L1Md elements identified as primary SPIN1 targets. Heatmap analysis across L1Md_A, L1Md_Gf, and L1Md_T elements revealed that H3K9me3 was clearly reduced in *Spin1*-KO cells (Fig 5C). The reduction in H3K9me3 was particularly evident at the 5′ end near the transcription start site, where SPIN1 binds most strongly. ATAC-seq heatmap analysis confirmed that chromatin accessibility increased at the same 5′ TSS-proximal regions in *Spin1*-KO cells across all three L1Md families (Fig 5D). These changes were specific to evolutionarily young families that carry the SPIN1-binding chromatin signature: the older LINE1 family L1Md_F and the ERVK family MMERVK10C, which lack H3K4me3 and are not bound by SPIN1, showed no detectable changes in either H3K9me3 or chromatin accessibility upon *Spin1* loss (Suppl Fig 4A,B). SPIN1-dependent chromatin regulation is therefore tightly coupled to the chromatin signature that defines its targets.

Given that SPIN1 cooperates with SPOCD1 and the DNMT3C/L machinery to direct DNA methylation of TEs in the male germline^17^, we asked whether SPIN1 also contributes to DNA methylation in mESCs. Bisulfite sequencing of the L1Md_T promoter, a young LINE1 element, revealed comparable CpG methylation in WT and *Spin1*-KO mESCs, with both genotypes showing similarly low methylation levels (Fig 5E). As a positive control, *Uhrf1*-KO mESCs (Suppl Fig 4C), which fail to maintain DNA methylation, showed near-complete loss of methylation at the same locus, confirming the assay’s sensitivity (Fig 5E; Suppl Fig 4D,E). The *Gapdh* promoter served as an unmethylated control. To extend this analysis to a different SPIN1 target TE class, we examined the IAPLTR1 promoter and likewise found no significant change in methylation between WT and *Spin1*-KO (Fig 5E; Suppl Fig 4D,E). DNA methylation at SPIN1 target loci is therefore unaffected by SPIN1 loss in mESCs, in contrast to the DNA methylation defects observed upon SPIN1 disruption in the germline^17^.

To identify H3K9 methyltransferases that may associate with SPIN1 and potentially contribute to H3K9me3 maintenance at TE loci, we examined candidate enzymes by targeted co-immunoprecipitation. SETDB1 was readily detected in Flag-SPIN1 immunoprecipitates but not in Flag-EGFP control, and G9a was similarly co-precipitated with Flag-SPIN1 (Fig 5F), establishing physical associations between SPIN1 and these two H3K9 methyltransferases. In contrast, SUV39H2, another major H3K9 methyltransferase, was not detected in Flag-SPIN1 immunoprecipitates (Fig 5F), suggesting that SPIN1 preferentially associates with specific H3K9 methyltransferases, including SETDB1 and G9a, rather than uniformly associating with all tested H3K9me3-depositing enzymes. Among these, SETDB1 is the principal H3K9 methyltransferase that establishes H3K9me3 at mammalian TEs^38^ making it a plausible candidate enzyme that could contribute to SPIN1-associated H3K9me3 maintenance at TE loci, with potential contribution from G9a as well.

Together, these analyses indicate that in mESCs, SPIN1 enforces TE silencing by maintaining H3K9me3-based heterochromatin and restricting chromatin accessibility, while DNA methylation is not engaged. The physical association of SPIN1 with SETDB1 and G9a suggests that these H3K9 methyltransferases may contribute to this chromatin-based repression. The molecular consequence of SPIN1 loss in mESCs is thus a chromatin-level shift in which young TEs retain their H3K4me3 mark, lose H3K9me3, and become accessible, providing the chromatin basis for the transcriptional derepression.

Integrating these findings, we propose that SPIN1 engages distinct silencing machineries depending on the cellular context (Fig 6). In non-germline cells, SPIN1 recognizes evolutionarily young TEs through combined H3K4me3 and H3K9me3 binding by Tudor 1 and Tudor 2, acts with SPINDOC through Tudor 1 and Tudor 3, and associates with SETDB1 and G9a to reinforce H3K9me3-based heterochromatin and restrict chromatin accessibility, with MORC4 potentially contributing to the chromatin accessibility output. In the germline, the same SPIN1 protein cooperates with SPOCD1 and the PIWI–piRNA pathway to direct DNMT3C/L-mediated DNA methylation^16,17^. SPIN1 thus exemplifies how a single histone reader can engage distinct silencing effectors and produce different chromatin outputs at TEs depending on cellular context.

**Figure 6.**
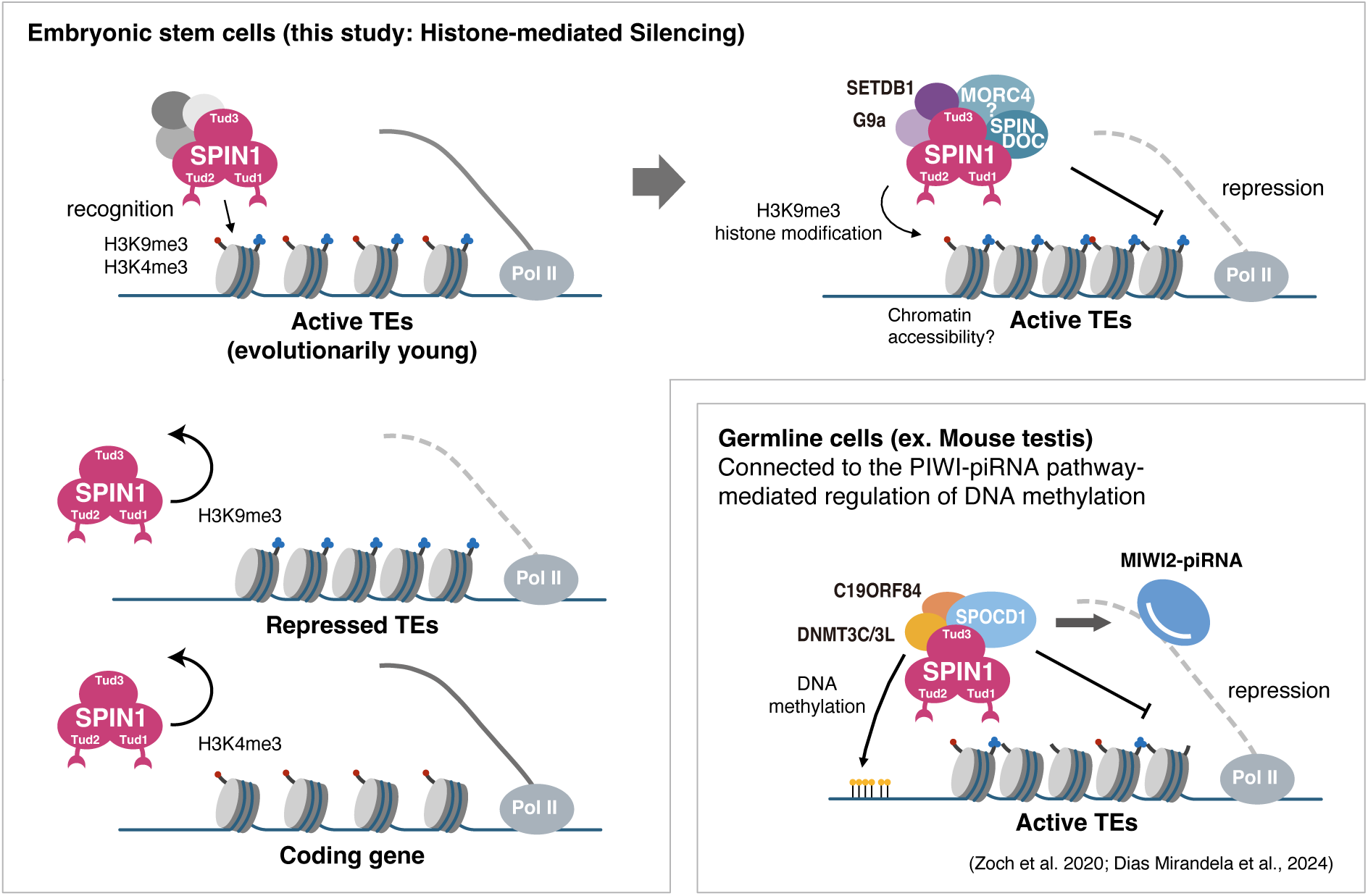
SPIN1 engages distinct transposon silencing machineries in germline and non-germline cells. A model for SPIN1-mediated transposon silencing across cellular contexts. In embryonic stem cells (top, this study), SPIN1 specifically recognizes evolutionarily young transposons carrying both H3K4me3 and H3K9me3 marks through its Tudor 1 and Tudor 2 domains. Notably, SPIN1 fails to bind loci marked solely by H3K9me3 (repressed TEs) or H3K4me3 (coding genes) (bottom left). Upon bivalent recognition, SPIN1 recruits the cofactor SPINDOC together with the H3K9 methyltransferases SETDB1 and G9a through Tudor 1 and Tudor 3. This complex reinforces H3K9me3-based heterochromatin and restricts chromatin accessibility, with MORC4 potentially contributing to the chromatin accessibility output (indicated by ‘?’). In the germline (bottom right, based on Zoch et al. 2020 and Dias Mirandela et al. 2024), SPIN1 instead cooperates with SPOCD1, the PIWI protein MIWI2, and the PIWI–piRNA pathway to direct DNMT3C/3L and C19ORF84 to deposit DNA methylation at piRNA-targeted TE loci. In both contexts, SPIN1 functions through its three Tudor domains but engages different cofactors and produces different chromatin outputs, illustrating how a single histone reader can be deployed in distinct silencing strategies adapted to different biological imperatives.

## Discussion

In this study, we identify SPIN1 as a chromatin-based effector of TE silencing in mESCs, where the PIWI–piRNA pathway is not operational. SPIN1 reads the combined H3K4me3/H3K9me3 chromatin signature of evolutionarily young TEs and engages the cofactor SPINDOC to enforce H3K9me3-based silencing and restrict chromatin accessibility without altering DNA methylation. SPIN1 also physically associates with the H3K9 methyltransferases SETDB1 and G9a, which represent candidate writers that may contribute to the H3K9me3 maintained at SPIN1 target loci. This mechanism is mechanistically distinct from the germline mode in which SPIN1 cooperates with SPOCD1 and the PIWI–piRNA pathway to direct DNA methylation, positioning SPIN1 as a context-dependent regulator that engages distinct silencing machineries in germline and non-germline cells and defines a chromatin-state-based mode of TE recognition that complements the established sequence– and RNA-based recognition pathways.

Mammalian cells deploy multiple TE silencing pathways that distinguish their targets through different molecular features. KRAB zinc finger proteins recognize specific DNA sequences within retroviral elements and recruit TRIM28/KAP1 to deposit H3K9me3^8–11^; the HUSH complex senses long, intronless nascent transcripts that characterize TE expression^12–14^; and the PIWI–piRNA pathway in germ cells employs sequence-specific small RNA guides^15–19^. The SPIN1-based silencing pathway we describe here adds a fourth recognition layer in which the chromatin state itself, namely the simultaneous presence of H3K4me3 and H3K9me3, serves as the targeting signal. Rather than reading a feature intrinsic to the TE sequence or its transcript, SPIN1 reads a chromatin configuration that emerges when an evolutionarily young, transcription-permissive TE acquires repressive marks. In this context, H3K4me3 likely reflects retained transcription potential at intact young TE promoters, whereas H3K9me3 most plausibly reflects ongoing silencing attempts by upstream pathways such as KRAB-ZFP/TRIM28 and HUSH. The simultaneous presence of both marks therefore identifies loci where transcriptional activity and silencing attempts are in active opposition, and SPIN1 functions as a sensor for incomplete silencing that consolidates repression by maintaining the H3K9me3-based heterochromatin state. KRAB-ZFPs and HUSH both recruit SETDB1 to establish H3K9me3 at TEs, suggesting that SPIN1 may stabilize their products by reading the resulting H3K4me3/H3K9me3 state and reinforcing the repression, providing robustness to the silencing network at loci where transcriptional escape is most likely. The recognition strategy of SPIN1 also stands out among other H3K9me3 readers. TNRC18, for example, recognizes H3K9me3 through its C-terminal BAH domain and silences ERV class I elements such as LTR12^39^, but its specificity for a particular ERV class implies that additional features beyond H3K9me3 alone constrain its targeting. By contrast, SPIN1 uses the combined H3K4me3/H3K9me3 chromatin signature as a major determinant of target selection, binding even to host genes that carry both marks (Suppl Fig 2A); this targeting strategy nonetheless selectively concentrates SPIN1 at transcription-permissive young TEs, where transcriptional escape would be most damaging.

The contrast between the germline and non-germline modes of SPIN1 action illuminates how the same chromatin reader can be repurposed for different silencing strategies. In the germline, SPIN1 binds SPOCD1 through Tudor 3 and contributes to piRNA-directed DNMT3C-mediated de novo DNA methylation of young L1 elements^16,17^, providing a stable, heritable layer of TE silencing that persists through global epigenetic reprogramming in the early embryo, an essential requirement for ensuring genome integrity across generations. The germline and non-germline modes share the common feature of using Tudor 1 and Tudor 3 to engage cofactors, but differ in the identity of the cofactor, the targeting signal, and the downstream chromatin output. This division reflects the different biological imperatives of the two contexts, namely transgenerational inheritance of silencing in the germline and within-lifespan maintenance of silencing in somatic and pluripotent cells. SPIN1 thus exemplifies how a single histone reader can acquire context-specific functions through partner-switching, enabling integrated control of TEs across germline and somatic compartments.

Our findings establish that SPIN1 is required to maintain H3K9me3 at its target TE loci (Fig 5A) and that SPIN1 physically associates with the H3K9 methyltransferases SETDB1 and G9a (Fig 5F). However, the precise mechanism by which SPIN1 contributes to H3K9me3 maintenance remains to be determined. Two non-exclusive scenarios are consistent with our data. First, SPIN1 may stabilize existing H3K9me3 deposited by SETDB1 or G9a, for example by retaining these methyltransferases at the locus to counter active H3K9 demethylation or to prevent displacement of methylated nucleosomes. Second, SPIN1 may actively recruit SETDB1 and/or G9a to introduce H3K9me3 de novo at TE loci that have acquired the initial H3K4me3/H3K9me3 chromatin signature, thereby reinforcing the silencing state established by upstream pathways such as KRAB-ZFP/TRIM28 or HUSH. Discriminating between these models will require time-resolved measurements following acute SPIN1 perturbation and direct tests of whether SETDB1 and G9a chromatin occupancy at SPIN1 target sites depends on SPIN1.

Our IP-MS analysis identified MORC4 alongside SPINDOC as a Tudor 1 and 3-dependent SPIN1 interactor, raising the possibility that MORC4 might also participate in SPIN1-associated chromatin regulation. The MORC family comprises GHKL-type ATPases that have emerged as important regulators of chromatin compaction and silencing. Notably, MORC2 functions as part of the HUSH complex to silence young LINE1 elements through ATP-dependent compaction of chromatin^40,41^, indicating that MORC family ATPases can directly contribute to the establishment of a closed chromatin state. By analogy, it is conceivable that MORC4 could similarly contribute to chromatin compaction at SPIN1 target TE loci, although this remains speculative in the absence of functional data. The reduced H3K9me3 and increased chromatin accessibility observed in *Spin1*-KO cells need not be linked through a single mechanism. The increased accessibility could arise simply as a passive consequence of reduced H3K9me3, or it could additionally involve the loss of an active compaction activity, potentially mediated by MORC4 in parallel with H3K9me3 maintenance, and these two possibilities are not mutually exclusive. Direct functional analysis of MORC4 in this context will be required to test these hypotheses.

Although our domain mutant analysis focused on H3K4me3 and H3K9me3 recognition, prior structural and biochemical studies indicate that SPIN1 has a broader histone modification readout capability. The Tudor 1/Tudor 2 binding pocket of SPIN1 can engage trimethylated lysines on histone H3 and H4 with varying affinities, and the combination of H3K4me3 with H3R8me2a or H3K9me3/2 has been reported to enhance binding through multivalent interactions^20–23^. These observations raise the possibility that SPIN1 may function as a multivalent reader whose recruitment is potentiated by the density and combination of multiple methyl marks at a given locus, rather than by a single mark in isolation. This model would explain why SPIN1 is enriched at H3K4me3-positive young L1Md and IAPEz copies but not at older TE families that carry H3K9me3 alone. Future work using SPIN1 structures with mutated reader pockets and quantitative binding assays with combinatorially modified peptides will be needed to test whether multivalent histone modification recognition contributes to SPIN1 target selectivity beyond the H3K4me3 and H3K9me3 modifications examined here.

Beyond the germline versus non-germline contrast discussed above, SPIN1 illustrates context-dependent switching of its functional output within non-germline cells themselves, as exemplified by its dual role as a repressor at TE loci and as a previously reported activator at rDNA promoters, where SPIN1 binding promotes chromatin opening and stimulates RNA polymerase I-mediated rRNA transcription^20,23^. Three non-exclusive factors likely contribute to these opposite outputs. First, although SPINDOC is implicated in both contexts, the surrounding effector machinery differs, with Pol I-associated factors at rDNA and MORC4 together with H3K9 methyltransferases at TE loci. Second, the local chromatin environment differs, with an H3K4me3-dominant permissive context at rDNA versus a dual H3K4me3/H3K9me3 heterochromatin-prone context at TE loci. Third, the polymerase machinery itself differs, Pol I at rDNA versus Pol II at TEs. Together with the germline mode discussed earlier, these contrasts establish SPIN1 as a context-sensitive scaffold whose interactome and output are dictated by the cellular context, the local chromatin landscape, and the polymerase machinery present at the bound locus, a versatility that may be a general feature of histone readers that integrate multiple inputs into context-appropriate outputs.

The physiological significance of SPIN1-mediated TE silencing in non-germline contexts remains to be fully established. *Spin1*-KO mice show postnatal lethality and skeletal muscle defects^27^, and *Spin1* is broadly expressed across embryonic and adult tissues^26^, yet the extent to which TE derepression contributes to these phenotypes is unclear. Given the connections between aberrant TE activation and pathologies including cancer, neurodegeneration, and autoimmunity^3,42–45^, our findings suggest that loss of SPIN1 function could contribute to disease through TE reactivation in addition to the rRNA synthesis and Wnt/β-catenin signaling defects previously described. The reliance on histone-state-based, rather than DNA-methylation-coupled, TE silencing may be particularly important in cellular states characterized by global DNA hypomethylation, such as mouse embryonic stem cells, preimplantation embryos, and primordial germ cells during epigenetic reprogramming^46,47^. In these contexts where DNA-methylation-coupled silencing is partially attenuated, chromatin-state-based silencing by SPIN1 may provide an essential compensatory layer of TE control, a hypothesis that could be tested by stage-specific *Spin1* perturbations during early development.

In conclusion, our study identifies SPIN1 as a chromatin-state reader that adds a histone-modification-based layer to the existing repertoire of mammalian TE silencing mechanisms, providing a piRNA-independent mechanism for selective TE silencing in non-germline cells that complements the DNA-methylation-coupled mechanism deployed in the germline. These findings expand our understanding of how the host genome integrates multiple recognition modes to constrain TE activity, and raise new questions about the cofactors, multivalent histone modification readout, and physiological contexts in which this chromatin-based silencing operates.

## Methods

### Plasmid construction

The pCAGGS-TetR-PA and pCAGGS-3×Flag vectors were derived from a pCAGGS backbone kindly provided by Dr. Hirotsugu Ishizu. To construct the pCAGGS-TetR-PA-mScarlet vector, a synthesized λN-PA cassette (Integrated DNA Technologies) was first inserted into this backbone. Subsequently, the λN sequence was replaced with a TetR coding sequence PCR-amplified from a Tet-ON plasmid kindly provided by Dr. Youjia Guo. To generate expression vectors for Flag-SPIN1, Flag-SPINDOC, and Flag-EGFP, their respective cDNAs (with the EGFP fragment PCR-amplified from another vector) were individually cloned into the pCAGGS-3×Flag vector using NEBuilder HiFi DNA Assembly Master Mix (New England Biolabs). Similarly, the PA-SPIN1 vector was constructed by inserting the *Spin1* cDNA into pCAGGS-TetR-PA. DNA fragments of *Spin1* mutants (ΔIDR, ΔTud1, ΔTud2, ΔTud3, Y98R, F141A, and Y170A) were PCR-amplified and subcloned into the restriction-digested pCAGGS-3×Flag or pCAGGS-TetR-PA vectors.

The pTIGRE-TetO-IAPEz-12MS2-ZsGreen-DR reporter construct was assembled using a multi-step cloning strategy. First, the *TIGRE* locus sequence was PCR-amplified from mESC genomic DNA and cloned into a pBluescript vector to serve as the backbone. In parallel, a synthesized ZsGreen coding sequence (Integrated DNA Technologies) was cloned into a pEBNA vector kindly provided by Dr. Kensaku Murano. Upstream TetO elements and an EF1α promoter were then assembled to create a TetO-EF1α-BoxB-ZsGreen cassette. The IAPEz sequence, amplified from mESC genomic DNA, was subsequently introduced via standard restriction enzyme cloning. This entire reporter cassette was then inserted into an endogenous NheI site within the cloned *TIGRE* sequence, yielding the intermediate construct pTIGRE-TetO-IAPEz-10×BoxB-ZsGreen. Finally, the BoxB sequence was replaced with 12×MS2 stem-loops, and the DR element was introduced.

All cloning steps were performed using NEBuilder HiFi DNA Assembly Master Mix or standard restriction enzyme-based methods. For the generation of *Pcgf6*-, *Spin1*-, *Uhrf1*-, and *Spindoc*-KO cells, as well as for targeted integration into the *TIGRE* locus, the respective single guide RNAs (sgRNAs) were cloned into the pSpCas9(BB)-2A-Puro (PX459) vector (Addgene, Plasmid #62988) by annealing synthesized oligos and ligating them into the vector. All PCR primers and sgRNA sequences are listed in Supplementary Table S1.

### Cell culture

Mouse embryonic stem (mES) cells were maintained on 0.1% gelatin-coated dishes at 37 °C in a humidified incubator with 5% CO_2_. The cells were cultured in Dulbecco’s Modified Eagle Medium (DMEM; 4.5 g/L glucose, with L-glutamine, without sodium pyruvate; Nacalai Tesque) supplemented with 10% heat-inactivated fetal bovine serum (FBS; Gibco), 1× MEM non-essential amino acids (Gibco), 1× GlutaMAX™ (Gibco), 0.2 mM sodium pyruvate (Gibco), 100 U/mL penicillin-streptomycin (Gibco), and 0.1 mM 2-mercaptoethanol (Gibco). To maintain the undifferentiated state, the medium was further supplemented with 1,000 U/mL recombinant mouse leukemia inhibitory factor (LIF; MBL), 1 μM PD0325901 (Fujifilm Wako), and 10 μM Y27632 (Fujifilm Wako).

### Construction of knockout mESCs

1.5 × 10^6^ mESCs were transfected with 3 μg of sgRNA expression plasmids using Lipofectamine 3000 Transfection Reagent (ThermoFisher Scientific) in a 6-well culture plate. At 24 h post-transfection, the cells were selected with 1 μg/mL puromycin at 37 °C for 72 h. The selection medium was then replaced with fresh medium containing 1 μg/mL puromycin for few days, and cells were further maintained to allow for colony formation and expansion.

### Tethering assay and flow cytometry

Tethering assay was performed as described previously^48,49^. mESCs (5 × 10^5^ cells/well) were seeded in 12-well plates and co-transfected with 2 μg of the pTIGRE-TetO-IAPEz-12MS2-ZsGreen-DR reporter plasmid and 1 μg of pCAGGS-TetR-fusion plasmids using Lipofectamine 3000 (Thermo Fisher Scientific). Doxycycline (Dox; 1 μg/mL) was added at the time of transfection. At 48 h post-transfection, the medium was replaced with fresh medium containing 1 μg/mL Dox for another 48 h. For flow cytometric analysis, cells were washed twice with PBS, detached using TrypLE Express (Thermo Fisher Scientific), and resuspended in PBS. The cell suspension was filtered through a 35-μm mesh-cap FACS tube (Corning). Data were acquired on a BD LSRFortessa X-20 cell analyzer (BD Biosciences). Color compensation was performed using single-color controls (cells transfected with either ZsGreen or mScarlet only) to account for spectral overlap. The gating strategy was as follows: (1) identification of the viable cell population based on FSC-A and SSC-A, (2) doublet discrimination using FSC-H/W and SSC-H/W parameters, and (3) selection of transfected cells based on mScarlet positivity (PE intensity > 10^3^) (Supplemental Fig1F). Within this mScarlet-positive population, the fluorescence intensity of ZsGreen (FITC channel) was analyzed to evaluate transcriptional repression. Data were processed using FlowJo software (BD Biosciences).

### RNA-seq analysis

Sequencing was performed with paired-end reads by HiSeq X or NovaSeq6000. Total RNA was extracted from control and *Spin1*-KO *Pcgf6*/*Dux*-dKO mESCs, as well as from wild-type and *Spin1*-KO mESCs, using ISOGEN II (Nippon Gene), and sequencing libraries were prepared by Macrogen Japan. Sequencing of the libraries was also performed by Macrogen Japan with 100 bp paired-end reads by NovaSeq6000. Paired– end sequencing reads were processed using the nf-core/rnaseq pipeline (v3.21.0)^50^. The pipeline was run under a Docker profile against the GRCm38 mouse reference genome. For gene-level expression analysis, transcript abundances were estimated using salmon as integrated within the nf-core/rnaseq pipeline^51^. Transcript-level estimates were imported and summarized to the gene level using tximport^52^, and differentially expressed genes (DEGs) were identified using DESeq2^53^ with default parameters. For TE expression analysis, the same raw sequencing reads were re-processed through the nf-core/rnaseq pipeline configured for repetitive-element quantification. Reads were aligned using STAR^54^ within the star_salmon mode of the pipeline, with the multi-mapping parameter set to permit up to 100 alignments per read (winAnchorMultimapNmax = 100, outFilterMultimapNmax = 100) to allow recovery of reads mapping to repetitive sequences. Pseudo-alignment was skipped (--skip_pseudo_alignment) and the minimum mapped reads threshold was removed (--min_mapped_reads 0). TE quantification was performed with TEcount from the TEtranscripts suite^55^, using gene annotations from the mm10 UCSC GTF and TE annotations from the TEtranscripts-provided mm10 repeat masker TE GTF (mm10_rmsk_TE.gtf). The –-stranded reverse option was applied based on the inferred library strandedness. TEcount aggregates reads at the level of TE subfamilies, allowing quantification of repetitive elements that cannot be uniquely assigned to individual genomic copies. Differential expression analysis of TE subfamilies was performed using edgeR^56,57^ on the TEcount output matrix. Volcano plots, heatmaps of selected TE subfamilies, and class-stratified boxplots (LINE, LTR, SINE, DNA) were generated in R using custom scripts.

### ATAC-seq Library Construction

ATAC-seq was performed as described previously^58,59^ with the following modifications. A total of 50,000 cells were washed with cold PBS and resuspended in 50 μL of cold lysis buffer containing 10 mM Tris-HCl (pH 7.5), 10 mM NaCl, 3 mM MgCl_2_, 0.1% NP-40, 0.1% Tween-20, and 0.01% digitonin. After incubation on ice for 3 min, the cells were washed with 1 mL of wash buffer containing 0.1% Tween-20 and centrifuged at 500 x *g* for 10 min at 4 °C to collect the nuclei. The nuclei pellet was then subjected to tagmentation in a 50 μL reaction mixture consisting of 2× tagmentation buffer (20% DMF), 0.01% digitonin, and Tn5 adapter complex (Diagenode). The reaction was incubated at 37 °C for 30 min with shaking at 1,000 rpm. Tagmented DNA was purified using the QIAGEN PCR purification kit and eluted in 23 μL of 0.1 × TE buffer. For library amplification, the purified DNA was mixed with Q5 Hot Start High-Fidelity 2× Master Mix (New England Biolabs) and uniquely barcoded i7 and i5 primers. PCR was performed with the following parameters: 72 °C for 5 min (gap filling), 98 °C for 30 s, followed by 18 cycles of 98 °C for 10 s, 63 °C for 15 s, and 72 °C for 1 min. The amplified libraries were purified and size-selected using 0.9 volumes of SPRIselect beads (Beckman Coulter). Library quality and quantity were assessed using a TapeStation (Agilent Technologies) and Qubit Fluorometer (Thermo Fisher Scientific).

### ATAC-seq analysis

Sequencing was performed with paired-end reads by NextSeq2000. Paired-end sequencing reads were processed using custom Snakemake workflows configured to retain multi-mapping reads for the analysis of repetitive elements. Reads were trimmed and aligned to the mouse GRCm38 (mm10) reference genome using Bowtie2^60^ with default parameters, retaining alignments to multiple genomic locations to enable analysis at TEs. Both primary alignments and unique alignments were retained for downstream analyses. PCR duplicates were removed using Picard MarkDuplicates to generate deduplicated BAM files. For analyses at TEs, primary alignment files were used to allow quantification of reads mapping to repetitive sequences. Normalized bigWig coverage tracks were generated with deepTools bamCoverage v3.5.0^61^ using –-binSize 1, –-normalizeUsing CPM, –-exactScaling, and –-ignoreForNormalization MT. Peak calling was performed using MACS2 callpeak^62^ on individual replicates and on merged BAM files. Average profile plots and heatmaps of chromatin accessibility were generated using deepTools computeMatrix together with plotProfile or plotHeatmap, centered either on SPIN1 CUT&Tag peak summits or across annotated TE elements stratified by family and by evolutionary age as described for CUT&Tag analyses.

### CUT&Tag Library Construction

CUT&Tag was performed as described previously^63^ with several modifications to optimize for mESCs. Briefly, 50,000 mESCs were lightly cross-linked with a final concentration of 0.1% formaldehyde in PBS for 2 min at room temperature, and the reaction was immediately quenched by adding glycine. After washing with 10% FBS/PBS, the cells were suspended in wash buffer [20 mM HEPES (pH 7.5), 150 mM NaCl, 0.5 mM spermidine, and 1x Protease inhibitor (Roche)] and immobilized on activated Concanavalin A-coated magnetic beads (Epicypher) in binding buffer [20 mM HEPES (pH 7.5), 10 mM KCl, 1 mM CaCl2, and 1 mM MnCl2]. The bead-bound cells were incubated with primary antibodies (listed in Supplementary Table S2) in 100 μL of antibody buffer [20 mM HEPES (pH 7.5), 150 mM NaCl, 0.5 mM spermidine, 0.01% digitonin, 2 mM EDTA, 0.1% BSA, and 1x Protease inhibitor (Roche)] overnight at 4 °C. Subsequently, cells were incubated with secondary bridge antibodies—either anti-rabbit IgG (H&L) (Rockland) or anti-mouse IgG (H&L) (Abcam)—diluted 1:100 in dig-wash buffer [20 mM HEPES (pH 7.5), 150 mM NaCl, 0.5 mM spermidine, 0.01% digitonin, and 1x Protease inhibitor (Roche)] for 1 h at room temperature. After washing, cells were incubated with pAG-Tn5 (Cell Signaling Technology) in dig-300 buffer [20 mM HEPES (pH 7.5), 300 mM NaCl, 0.5 mM spermidine, 0.01% digitonin, and 1x Protease inhibitor (Roche)] for 1 h at room temperature. Tagmentation was initiated by adding 10 mM MgCl_2_ and performed at 37 °C for 1 h. To stop tagmentation and reverse form formaldehyde crosslinks, 0.5 M EDTA, 10% SDS, and Proteinase K were added to each sample, followed by incubation at 67°C overnight. The de-crosslinked DNA fragments were then extracted using the phenol-chloroform method. The extracted DNA was amplified using NEBNext High-Fidelity 2× PCR Master Mix (New England Biolabs) with the following cycles: 72 °C for 5 min, 98 °C for 30 s, followed by 14 cycles of 98 °C for 10 s and 63 °C for 10 s. The resulting libraries were purified and size-selected using 0.9 volumes of SPRIselect beads (Beckman Coulter). Library quality was assessed using a TapeStation (Agilent Technologies).

### CUT&Tag analysis

Sequencing was performed with paired-end reads on a NextSeq2000. Paired-end sequencing reads were processed using the nf-core/cutandrun pipeline (v3.2.2)^50^ for read trimming, alignment to the mouse GRCm38 (mm10) reference genome with Bowtie2^60^, and quality filtering. The pipeline was run under a Docker profile with the following parameters: –-normalisation_mode CPM, –-normalisation_binsize 1000, –-minimum_alignment_q_score 0, –-trim_nextseq 20, and the ENCODE GRCm38 blacklist applied to exclude problematic genomic regions.

The downstream analysis followed the framework established by previous research^17^ with custom Snakemake workflows. Alignment files from the nf-core/cutandrun pipeline were processed to remove PCR duplicates using Picard MarkDuplicates or samtools markdup. Replicate BAM files were merged using samtools merge^64^ for downstream analyses. Normalized bigWig coverage tracks were generated using deepTools bamCoverage v3.5.0^61^ with the following parameters: –-binSize 1, –-normalizeUsing BPM, –-ignoreForNormalization MT. Log2 enrichment profiles of CUT&Tag samples over IgG controls were generated using deepTools bamCompare with the same parameters plus-e, –-scaleFactorsMethod None.

Peak calling was performed using MACS2 callpeak^62^ on individual replicates as well as on merged BAM files, with the corresponding IgG sample set as the control. The parameter –-keep-dup all was used to include duplicate reads when present. To obtain high-confidence peaks, peaks were filtered for a minimum coverage of 20 reads in the CUT&Tag sample and a peak score greater than the mean peak score across all called peaks. Peak overlaps between H3K4me3, H3K9me3, and SPIN1 were analyzed using BEDTools intersect^65^ in R.

Genomic feature annotation of peaks was performed in R using the GenomicRanges and GenomicFeatures packages^66^ together with the GRCm38 RepeatMasker annotation and Ensembl gene annotation. Each peak was assigned to a single genomic feature category through a hierarchical overlap analysis, in which peaks were classified sequentially in the following priority order: (1) LINE elements, peaks overlapping repeat annotations of the LINE class; (2) Other TE/repeat elements – genic, peaks overlapping LTR, Simple_repeat, Satellite, ERVK, Retrotransposon, or SINE annotations together with a gene unit; (3) Other TE/repeat elements – intergenic, peaks overlapping the same repeat annotations but outside any gene unit; (4) Genes, peaks overlapping a transcriptional unit but not any repeat; (5) Intergenic, peaks with no overlap to any of the features above. Each peak was thereby assigned exclusively to the highest-priority matching category.

Average profile plots and heatmaps centered on peak summits or across annotated TE elements were generated with deepTools computeMatrix and plotProfile or plotHeatmap using log2(IP/IgG) bigWig tracks. For analyses at TE subfamilies, coordinates were extracted from the RepeatMasker annotation, with size filtering applied where indicated. Stratification of L1Md_A elements by evolutionary age was based on sequence divergence from the consensus, as described^31^. For analyses stratifying TE elements by H3K4me3 status (Fig 2E), individual TE copies were clustered by k-means clustering (k = 2) on H3K4me3 signal intensity using deepTools plotHeatmap, separating them into H3K4me3-high and H3K4me3-low clusters.

### Rescue assay

A total of 2 × 10^6^ *Spin1*-KO cells were transfected with 3μg of Flag-SPIN1-HygR plasmid using Lipofectamine 3000 Transfection Reagent (ThermoFisher Scientific). At 24 h post-transfection, the cells were selected with 800 μg/mL hygromycin at 37 °C for 48 h. The selection medium was then replaced with fresh medium containing 800 μg/mL hygromycin for an additional 72 h. Following selection, the cells were harvested for RT-qPCR analysis (described below).

### RT-qPCR

Total RNA extraction and reverse transcription were performed as previously described^67^. Briefly, total RNA was extracted from mESCs using ISOGEN II (Nippon Gene). The extracted RNA was reverse transcribed into cDNA using the ReverTra Ace® qPCR RT Master Mix with gDNA Remover (Toyobo). Quantitative PCR (qPCR) was performed with TB Green® Premix Ex Taq™ (Tli RNaseH Plus) (Takara Bio) on a LightCycler® 96 System (Roche). The primer sequences used are listed in Supplementary Table S1. Relative mRNA expression levels were calculated using the 2^-ΔΔCt^ method, normalized to *Gapdh* expression. All data were obtained from three independent experiments.

### Co-immunoprecipitation

For most Co-IP experiments involving PA-tagged or Flag-tagged proteins, mESCs were first mildly cross-linked *in situ* to stabilize protein–protein interactions. Briefly, cells were incubated with 0.1% formaldehyde in PBS for 5 min at room temperature, followed by quenching with 100 mM glycine-NaOH (pH 7.5) for 4 min at room temperature. After washing with ice-cold PBS, cells were harvested and lysed in HEPES-RIPA buffer [20 mM HEPES-NaOH (pH 7.5), 150 mM NaCl, 1 mM MgCl_2_, 1 mM EGTA, 1% NP-40, 0.25% sodium deoxycholate, 0.05% SDS, 5 mM dithiothreitol (DTT), 0.5% Triton X-100, 1× cOmplete™, Mini, EDTA-free Protease Inhibitor Cocktail (Roche), and 1× PhosSTOP™], according to a previously described protocol with minor modifications (Nishino and Kosako 2022). The lysates were frozen at –80°C for at least 1.5 h, thawed, and homogenized using a Bioruptor II (Diagenode; Low power, 30 s ON / 30 s OFF for 5 cycles). The homogenized lysates were cleared by centrifugation at 10,000 × *g* for 10 min at 4°C.

For the native Co-IP experiment validating the SPIN1–SPINDOC interaction (Fig. 4C), the cross-linking, quenching, freezing, and sonication steps were omitted. Instead, cells were directly harvested, resuspended in a standard RIPA buffer [50 mM Tris-HCl (pH 8.0), 150 mM NaCl, 1% Triton X-100, 0.5% sodium deoxycholate, and 1× cOmplete™, Mini, EDTA-free Protease Inhibitor Cocktail (Roche)], and incubated on ice for 10 min. The lysates were then cleared by centrifugation at 10,000 × *g* for 10 min at 4°C.

The resulting supernatants from either lysis method were incubated with MagCapture HP Anti-PA tag Antibody Magnetic Beads (Fujifilm Wako), anti-Flag M2 monoclonal antibody (Sigma-Aldrich), or anti-SPIN1 antibody pre-bound to Dynabeads Protein G (Thermo Fisher Scientific) for 2 h at 4°C. Details of all antibodies used in this study, including their sources and dilutions, are provided in Supplementary Table S2. The beads were washed three times with their respective lysis buffers (HEPES-RIPA or standard RIPA). For western blot analysis, bound proteins were eluted by boiling in SDS sample buffer, resolved by SDS-PAGE, and detected as described below. For liquid chromatography-mass spectrometry (LC-MS) analysis, the washed beads were subjected to further processing as described in the “Mass spectrometry (MS) sample preparation” section.

### Western blotting

Western blotting was primarily performed as described previously^67^. Proteins were resolved by SDS-PAGE and transferred onto PVDF membranes (Fujifilm Wako). The membranes were blocked with 5% skim milk in PBS and then incubated with primary antibodies, followed by incubation with appropriate HRP-conjugated secondary antibodies. Details of all antibodies used in this study, including their sources and dilutions, are provided in Supplementary Table S2. All antibodies were diluted in PBS containing 0.1% Tween-20 (PBS-T) and 1% skim milk. The membranes were washed extensively with PBS-T between each step. Signals were visualized using Clarity Western ECL Substrate (Bio-Rad) and captured with a ChemiDoc MP Imaging System (Bio-Rad).

### Mass spectrometry (MS) sample preparation

Immunoprecipitation eluates in 50 µL 0.1 M Tris-HCl (pH 6.8), 4% SDS, 23.6% Glycerol, 0.2 M DTT were prepared for MS proteomics with the SP4 method^68^. Briefly, glass spheres (Supelco, USA) were washed and adjusted to a concentration of 0.5 μg/μL in acetonitrile, and 200 μL was added to each eluate and mixed. Samples were centrifuged at 16,000 x *g* for 5 min at RT, and the supernatants were discarded. Samples were washed 3 times with 80% ethanol, with centrifugation at 16,000 x *g* for 3 min at RT between each wash. Supernatants were discarded. Then 250 µL 50 mM ammonium bicarbonate containing 5 mM Tris(2-carboxyethyl)phosphine hydrochloride (TCEP.HCl), 20 mM 2-chloroacetamide (CAA), 0.02% lauryl maltose neopentyl glycol (LMNG)^69^, 200 ng Lysyl Endopeptidase (Lys-C), and 200 ng trypsin was added, and the samples were incubated at 37 °C for 18 hr with shaking at 1,000 r.p.m.

Digested samples were centrifuged at 16,000 x *g* for 5 min at RT. The supernatants were taken and kept. The beads were washed once with 150 µL, and this was combined with the main supernatant fractions. The samples were acidified with trifluoroacetic acid (TFA) to 0.8% v/v (pH 2-3). Then they were desalted with poly(styrenedivinylbenzene) copolymer (SDB-XC) StageTips (Nikkyo Technos, Japan)^70^. The samples were eluted from SDB-XC StageTips with 20 µL 0.1% TFA, 20% water, 80% acetonitrile with centrifugation at 1,000 x *g* for 5 min at RT, and the samples were vacuum dried. The tryptic peptides were dissolved in 11 µL 0.1% formic acid, 3% acetonitrile, 97% water and 5 µL was injected for MS.

### MS data collection

The samples were measured with a Q Exactive Plus mass spectrometer, coupled to an EASY-nLC 1200 apparatus, and a Nanospray Flex ion source (Thermo Fisher Scientific). Data were collected with data-independent acquisition (DIA)^71^. Peptides were separated with a 100-µm I.D analytical column filled with 1.9-µm C18 particles (Reprosil, Dr. Maisch, Germany), for 20-cm filling length. Mobile phase A was 0.1% formic acid in water and mobile phase B was 0.1% formic acid, 20% water, 80% acetonitrile. Solvents were liquid chromatography (LC)/MS grade. The peptides were measured over 90-minutes: the LC gradient started with solvent A only for the first minute, then increased linearly from 5 to 40% solvent B from one to 64 minutes, then increased from 40 to 95% solvent B from 64 to 66 minutes, solvent B was maintained at 95% from 66 to 74 minutes, then the concentration decreased from 95 to 5% from 74 to 75 minutes, and for the final 5 minutes solvent B was maintained at 5%. The flow rate was 150 nL/min.

The following MS settings were maintained throughout data acquisition: positive mode; electrospray voltage, 2.0 kV; capillary temperature, 250 °C; S-lens RF level, 50.0; centroid data. For data acquisition cycles, one full MS scan was done from 495-745 m/z at 70,000 resolution, 3e6 AGC target, and 100 ms maximum IT. This was followed by 25 DIA scans with isolation windows of 10 m/z covering the precursor mass range of 495 – 745 m/z. A default charge state of 3, 3e6 AGC, and auto maximum IT were applied. Normalized HCD collision energies were 27%, and the fixed first mass was 200 m/z.

### MS data analyses

Raw data files were processed using DIA-NN software^72^, version 2.3.2, to produce a protein group matrix file listing the relative protein abundancies. A predicted spectral library was first made using DIA-NN with a *Mus musculus* Swiss-Prot reviewed and TrEMBL unreviewed FASTA file proteome database (downloaded from UniProt in August 2025). The contaminants FASTA file database provided by DIA-NN was also included in the library generation and search of the raw data files. Also, the additional option “--duplicate-proteins” was applied to generate the library and search the data, so that duplicate entries were not skipped. The default setting of the precursor false discovery rate (FDR) was 1%.

The resulting protein group matrix file was analyzed using Perseus software (version 2.1.1.0). Potential contaminants and reverse database hits were removed. Protein intensities were log2-transformed, and missing values were imputed with random numbers drawn from a normal distribution (width: 0.3, down shift: 1.8) to simulate the detection limit. Significant interactors were identified using a two-sample Student’s t-test (FDR < 0.05, S0 = 0.5) comparing the PA-SPIN1 group against the PA-mScarlet control group, the PA-SPIN1-ΔTud1 group, and the PA-SPIN1-ΔTud3 group, respectively (Supplementary Table S3).

For the Venn diagram, the bait protein SPIN1, immunoglobulin-derived entries, and ambiguous predicted protein groups were excluded as non-specific or non-informative hits. Specifically, immunoglobulin heavy/light chain variable regions (Ighv5-15), which were likely derived from antibody leakage during the elution process, were excluded. Additionally, uncharacterized protein clusters with unrefined annotations (including Gm20736;Gm20890;Gm20894;Gm20905;Gm21095;Gm21173;Gm21258;Gm21294;G m21627;Gm28553;Gm29564;Gm29866;Gm31571) were also omitted from the final interaction network to focus on characterized functional interactors.

### Bisulfite Sanger sequencing

Genomic DNA was extracted using the DNeasy Blood & Tissue Kit (Qiagen) according to the manufacturer’s instructions. Bisulfite conversion of the DNA was performed using the EZ DNA Methylation-Lightning Kit (Zymo Research). The targeted regions were amplified using TaKaRa EpiTaq HS (for bisulfite-treated DNA) (Takara Bio). Primers were designed using MethPrimer software^73^, and their sequences are listed in Supplementary Table S1. The PCR conditions for *Gapdh*, L1Md_T, and IAPLTR1 were as follows: 40 cycles of denaturation at 98 °C for 10 s, annealing for 30 s (at 57 °C for *Gapdh*, 59 °C for L1Md_T, and 53 °C for IAPLTR1), and extension at 72 °C for 30 s. The amplified products were gel-purified, subcloned into the T-Vector pMD20 (Takara Bio), and subjected to Sanger sequencing using the M13 reverse primer. At least 24 independent clones were sequenced per sample. The sequencing results were analyzed to evaluate methylation patterns using the QUMA software^74^.

## Acknowledgments

We thank Sefan Asamitsu, Yoshifumi Fujioka, and the other members of the Laboratory for Functional Non-coding Genomics at RIKEN IMS for technical assistance and helpful discussions. We are grateful to Haruhiko Siomi (Chiba University) and Yoichi Shinkai (RIKEN) for insightful discussions and for sharing materials. We thank Kurumi Shinoda for experimental assistance. This study was supported by JSPS KAKENHI grants 24H02060, 25H01305, 26K01941, The Sumitomo Foundation Grant (to Y.W.I.), and 24K18057 (to H.Y.). Y.W.I. is also supported by JST FOREST (JPMJFR224L). H.Y. is supported by the Japan Society for the Promotion of Science (25KJ0408) and the Takeda Science Foundation (2024047451).

## Author contributions

H.Y. and Y.W.I. conceived the study. H.Y. performed most experiments with C.T. who generated dKO cells and set up tethering reporter assay system. C.B. and K.I. performed mass spectrometry analyses. N.Y.-K. and H.K. contributed to chromatin analyses. A.S. contributed to experimental validation. H.Y. and Y.W.I. wrote the manuscript with input from all authors.

## Competing interests

The authors declare no competing interests.

## Supplementary Figure legends

**Figure S1.**
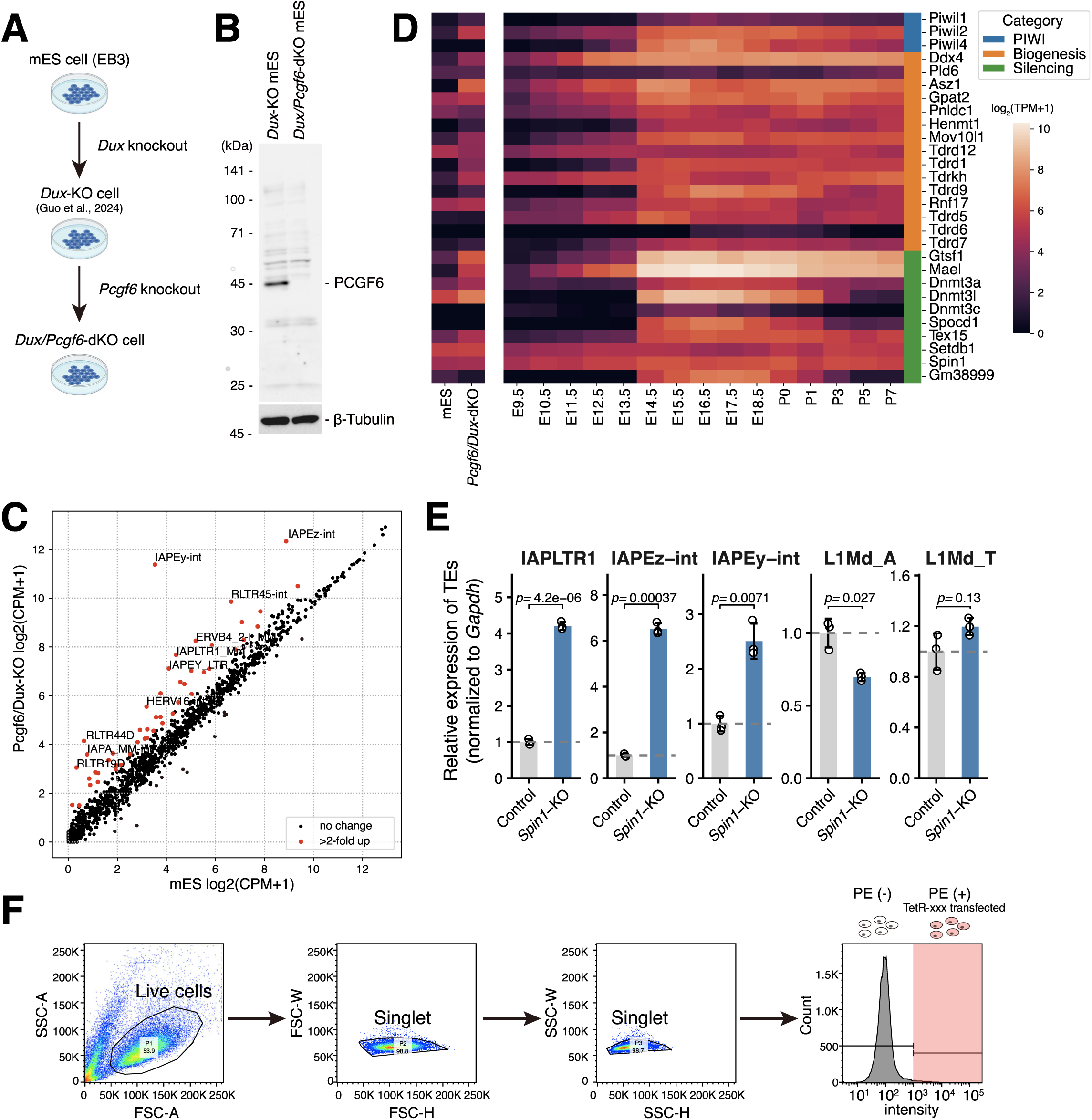
Generation of *Pcgf6/Dux*-dKO ESC system. (A) Schematic workflow for the generation of *Dux/Pcgf6*-dKO mESCs. Starting from EB3 mESCs, *Dux* was first deleted to prevent 2C-like arrest, followed by *Pcgf6* knockout to induce TE desilencing while maintaining cellular proliferation. (B) Western blot analysis confirming the loss of PCGF6 protein in *Pcgf6/Dux*-dKO ESCs. β-Tubulin was used as loading control. (C) Scatter plot comparing TE expression between WT and *Pcgf6/Dux*-dKO mESCs based on RNA-seq data. TEs significantly upregulated in dKO cells (log_2_ fold change ≥ 1) are highlighted in red. (D) Heatmap showing the expression of piRNA pathway factors (categorized into PIWI (blue), Biogenesis (orange), and Silencing factors (green)) in WT and *Pcgf6/Dux*-dKO mESCs compared to various stages of male germline development (embryonic days E9.5–E18.5 and postnatal days P0–P7). While certain factors like *Mael* and *Piwil2* show weak induction in dKO cells, key nuclear silencing factors such as *Piwil4 (Miwi2)* and *Spocd1* remain undetectable in *Pcgf6/Dux*-dKO mESCs. (F) RT-qPCR analysis of representative TEs (IAPLTR1, IAPEz-int, IAPEy-int, L1Md_A, L1Md_T) in WT and *Spin1*-KO ESCs. Expression levels were normalized to *Gapdh*. Data are presented as mean ± SEM from three independent experiments (*n* = 3). *P*-values were calculated using a two-sided Student’s t-test. (G) Gating strategy for flow cytometry analysis of the tethering assay. Sequential gates were applied to identify live cells (SSC-A vs. FSC-A) and single cells (FSC-W vs. FSC-H, followed by SSC-W vs. SSC-H). The ZsGreen intensity was then analyzed in the PE-positive population, representing cells successfully transfected with the TetR-fusion constructs.

**Figure S2.**
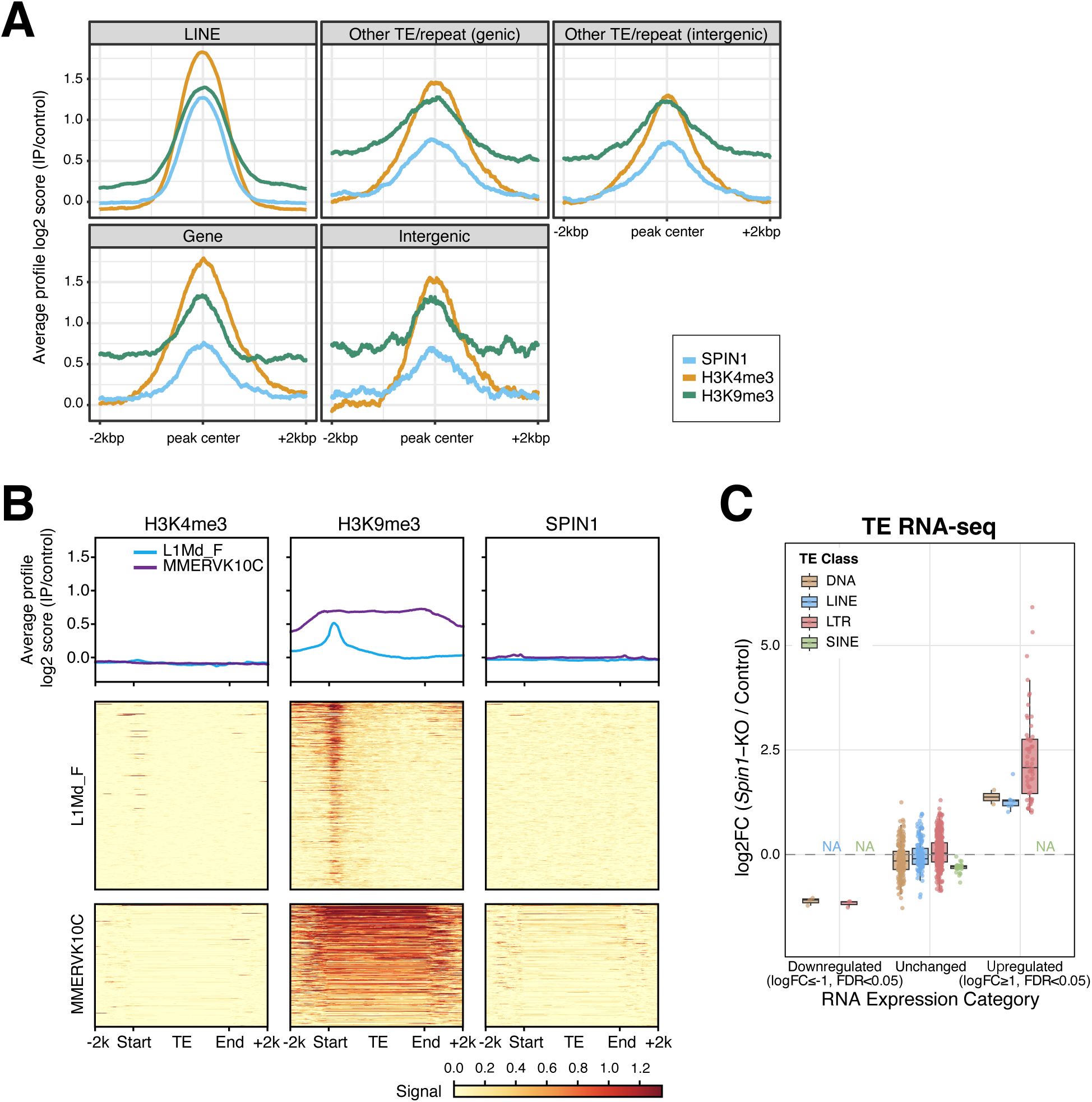
Genome-wide SPIN1 binding analyses. (A) Average CUT&Tag profiles of SPIN1 (light blue), H3K4me3 (orange), and H3K9me3 (green) centered on peak regions across various genomic categories, including LINEs, other TE/repeats (genic and intergenic), genes, and intergenic regions. (B) Average CUT&Tag profiles (top) and heatmaps (bottom) for H3K4me3, H3K9me3, and SPIN1 at L1Md_F and MMERVK10C families. These TE families exhibit H3K9me3 enrichment but lack H3K4me3 and SPIN1 binding, indicating that old inactive TEs with H3K9me3 alone are insufficient for SPIN1 recruitment. (C) Box plots showing the distribution of log_2_ fold changes in TE expression in *Spin1*-KO mESCs relative to control cells, categorized by TE class (DNA, LINE, LTR, and SINE) and regulation status. Upregulated TEs are defined by a log_2_ fold change ≥ 1 and FDR < 0.05, while downregulated TEs are defined by a log_2_ fold change ≤ –1, FDR < 0.05. LTR and LINE elements represent the majority of upregulated TEs, where LTR elements are highly upregulated.

**Figure S3.**
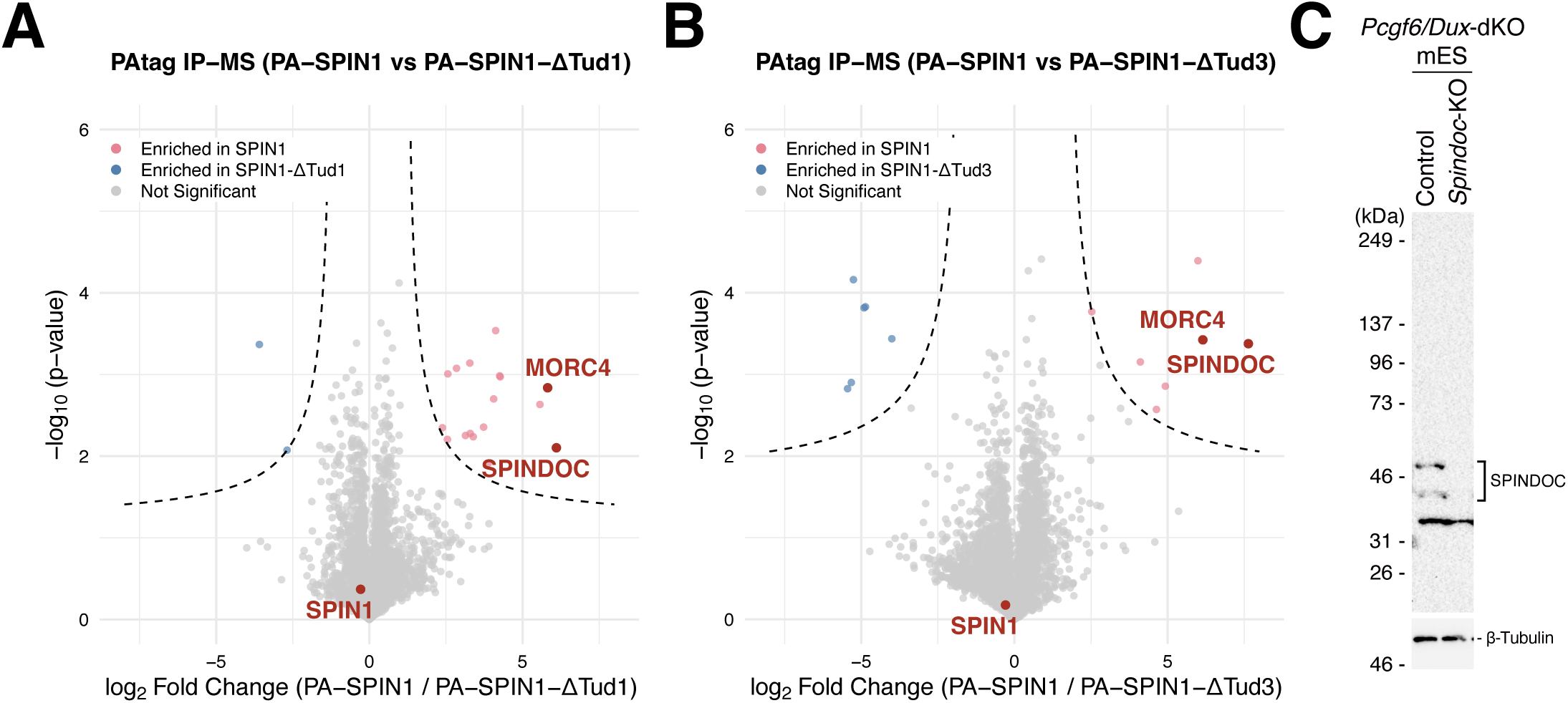
Proteome analysis of SPIN1 mutant interactors. (A), (B) Volcano plots showing the enrichment of proteins co-purified with PA-tagged full-length SPIN1 (PA-SPIN1) compared to (**A**) PA-SPIN1-ΔTud1 or (**B**) PA-SPIN1-ΔTud3, as determined by quantitative mass spectrometry (IP-MS). The dashed horizontal line denotes FDR = 0.05. Proteins significantly enriched in the WT control indicate Tud1– or Tud3-dependent interactors, respectively. SPINDOC and MORC4 are highlighted (A full list of identified proteins is provided in Supplementary Table S3). (C) Western blot analysis confirming the loss of SPINDOC protein in *Spindoc*-KO cells. β-Tubulin was used as loading control.

**Figure S4.**
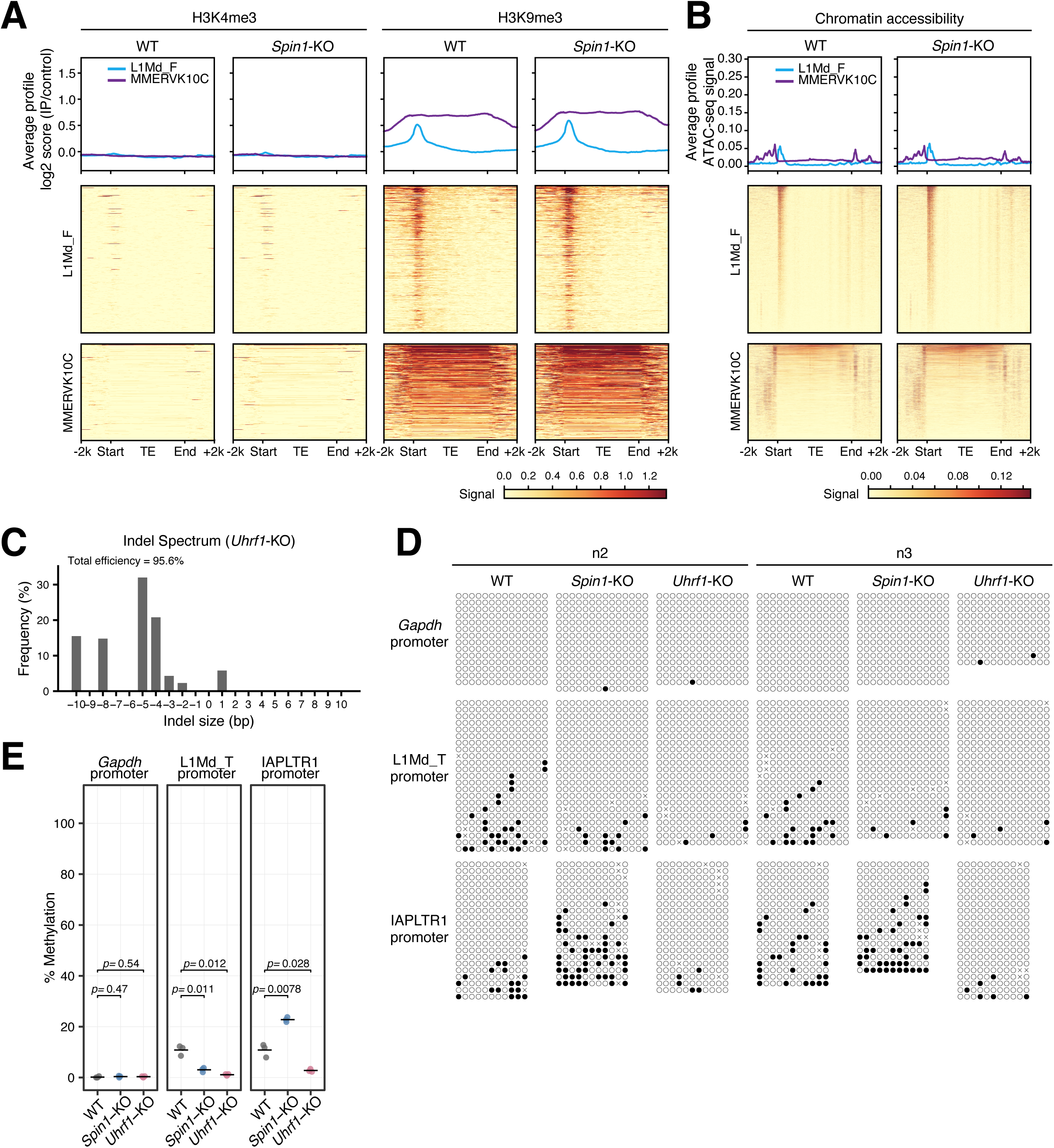
Chromatin and DNA methylation analyses. (A), (B) Average CUT&Tag profiles and heatmaps of H3K4me3 and H3K9me3 (**A**) and ATAC-seq (**B**) at TE families lacking SPIN1 binding (L1Md_F and MMERVK10C). Consistent with the lack of SPIN1 recruitment (see Supplementary Fig. 2b), H3K9me3 levels and chromatin accessibility remain unchanged in *Spin1*-KO cells at these loci. (C) Indel spectrum of *Uhrf1*-KO cells generated by CRISPR/Cas9. Genomic DNA from control (WT) and *Uhrf1*-KO cells was subjected to Sanger sequencing and subsequent TIDE (Tracking of Indels by Decomposition) analysis^75^. The bar graph displays the frequency (%) of insertions (+) and deletions (-) of various sizes (bp) at the targeted locus. The high frequency of frameshift mutations, particularly the prominent –5 bp and –4 bp deletions, confirms the successful disruption of the *Uhrf1* gene. (D) Bisulfite sequencing analysis of DNA methylation at the *Gapdh* promoter (control) the L1Md_T promoter, and the IAPLTR1 promoter in WT, *Spin1*-KO, and *Uhrf1*-KO cells. Filled and open circles represent methylated and unmethylated CpG sites, respectively. *Uhrf1*-KO serves as a positive control for the loss of DNA methylation. (E) Quantification of DNA methylation levels from the bisulfite sequencing data. Data are presented as mean ± SEM from three independent experiments (*n* = 3). *P*-values were calculated using a two-sided Student’s t-test.

## Supplementary Table legends

**Supplementary Table S1. Oligonucleotides and primers used in this study.**

This table lists the primers for plasmid construction, RT-qPCR and bisulfite sequencing analyses, and sequences of single guide RNAs (sgRNAs) for CRISPR/Cas9-mediated knockout.

**Supplementary Table S2. Antibodies used in this study**.

This table provides the details of primary and secondary antibodies used for Western blotting, immunoprecipitation, and CUT&Tag, including the target, host species, clonality, application-specific dilutions, company, and catalog number.

**Supplementary Table S3. Mass spectrometry analysis of SPIN1-interacting proteins**.

This table contains three sheets detailing the proteins identified by LC-MS/MS in the immunoprecipitates of PA-SPIN1, its Tudor domain deletion mutants (PA-SPIN1-ΔTud1 and PA-SPIN1-ΔTud3), and the PA-mScarlet control from mESCs. Sheet 1 (SPIN1 vs mScarlet), Sheet 2 (SPIN1 vs SPIN1-ΔTud1), and Sheet 3 (SPIN1 vs SPIN1-ΔTud3) provide protein and gene annotations, the number of total and proteotypic sequences (unique peptides) identified, and the log2-transformed intensities for three independent biological replicates per group. Statistical analysis was performed using a two-sample Student’s t-test in Perseus. “Log2 Fold Change” represents the difference in mean log2 intensities between the two groups compared in each sheet. “-Log10 p-value” and “q-value” indicate the statistical significance and permutation-based false discovery rate (FDR), respectively. Proteins that passed the significance threshold (FDR < 0.05, S0 = 0.5) are marked with a “+” in the “Significant” column.

## Notes

### Competing Interest Statement

The authors have declared no competing interest.

